# A systematic delineation of 3′UTR regulatory elements and their contextual associations

**DOI:** 10.1101/2025.06.09.658412

**Authors:** Nan Liang, Jing-Yi Li, Yang Ding, Yu Wang, Jackson Champer, Li-Chen Ren, Ge Gao

## Abstract

The 3′UTR encodes regulatory information that shapes transcript abundance and subcellular distribution, yet the underlying sequence rules remain incompletely defined. Using SEERS, we quantified the effects of ∼2 million synthetic 3′UTR inserts in A549 and HCT116 cells on RNA output and nuclear-cytoplasmic partitioning. We identify a broad repertoire of short (2∼8 nt) elements whose effects largely align along a major coupled axis that links high expression with cytoplasmic enrichment and low expression with nuclear enrichment, and contribute predominantly in an additive manner. A context-aware deep learning model (TALE) captures most of this behavior while revealing that strong context dependence is uncommon and emerges mainly in rare, extreme contexts consistent with higher-order constraints. Applying TALE to ClinVar 3′UTR variants prioritizes a subset of pathogenic SNVs with aberrant predicted effects, frequently explained by creation or disruption of splice-site-like U1 telescripting signals, highlighting a potent route to 3′UTR-driven disease mechanisms.

## INTRODUCTION

In human cells, the 3′ untranslated region (3′UTR) serves as a major post-transcriptional regulatory module: by shaping mRNA stability, translational efficiency, and nuclear-cytoplasmic partitioning and localization, it substantially expands the plasticity and complexity of gene regulatory networks.^1–4^ Unlike promoters and enhancers, which act primarily at the transcriptional level, 3′UTRs are thought to exert most of their control through post-transcriptional mechanisms, most notably by recruiting RNA-binding proteins (RBPs) and microRNA effector complexes that collectively determine mRNA fate.^3^ A number of representative cis-regulatory signals within 3′UTRs have been described, including miRNA binding sites, AU-rich elements (AREs), localization “zipcode” sequences, and motifs linked to 3′-end processing. Yet these dispersed rules have not converged into a coherent “grammar” explaining how short elements, in combination and in specific sequence contexts, jointly specify transcript abundance and subcellular localization.

A central obstacle to delineating 3′UTR grammar is confounding from the endogenous genomic environment. The output of a given gene reflects not only 3′UTR-encoded regulation, but also transcriptional inputs from promoters and distal enhancers, chromatin state, and three-dimensional genome organization, and it is coupled to co-transcriptional processes such as splicing, 3′-end formation, and alternative polyadenylation.^5–8^ Moreover, many genes produce multiple 3′UTR isoforms, further complicating efforts to attribute observed phenotypes cleanly to sequence features within a specific 3′UTR.

Inspired by paradigms developed for promoters and enhancers,^9–16^ massively parallel reporter assays (MPRAs) have been widely applied to study 3′UTR function.^17–21,2,22–31^ By embedding many candidate 3′UTR sequences into a common reporter transcript and cellular context, MPRA-based approaches can isolate 3′UTR-driven effects on RNA output and localization under controlled conditions. However, most existing 3′UTR MPRA studies rely on naturally occurring sequences with limited diversity, restricting coverage of the relevant sequence space.^32–34^ In addition, the coupling between transcript abundance and subcellular localization remains less systematically characterized.^3,25,35–37^ More broadly, recent MPRA work has highlighted that standard workflows can induce measurable cellular perturbations and stress responses, potentially enriching workflow-associated signals (e.g., innate immune activation) and introducing confounding into the readout.^38,39^ Together, these limitations impede comprehensive identification of 3′UTR regulatory elements and combinatorial rules, and they complicate principled modeling and visualization of context dependence and interactions.

In vitro-synthesized random sequences have proven powerful for revealing binding motifs and context preferences of RBPs and microRNAs.^40,41^ More generally, strategies that combine randomized sequence libraries, MPRA-style measurements, and machine learning have been notably successful in decoding in vivo regulatory programs governing RNA splicing, 5′UTRs, alternative polyadenylation, and promoters.^42–46^ Building on these advances, we reason that a more comprehensive and less biased understanding of 3′UTR regulatory elements and their contextual associations can be achieved by systematically measuring the effects of large numbers of random 3′UTR sequences on transcript abundance and nuclear-cytoplasmic distribution, and by learning the mapping from sequence to regulatory phenotypes using context-aware neural networks. This framework offers broad sequence-space coverage with fewer prior assumptions, and it provides a testable basis for interpreting the functional impact of disease-associated 3′UTR variants.

## RESULT

### SEERS reduces cellular perturbations associated with standard MPRA workflows

To mitigate cellular perturbations associated with conventional MPRA workflows, we developed SEERS (Selective Enrichment of Episomes with Random Sequences) (**Fig. 1A**). SEERS leverages an EBNA1/OriP-based episomal vector^47^ (**Supplemental Fig. S1**) that tethers to host chromosomes and replicates and segregates with the cell cycle, enabling long-term episome maintenance. This design allows puromycin-based selection and expansion of successfully transfected, surviving cells as they recover from transfection-associated toxicity, thereby reducing non-specific intracellular disturbances commonly observed in standard MPRA protocols.^38^

**Fig. 1.**
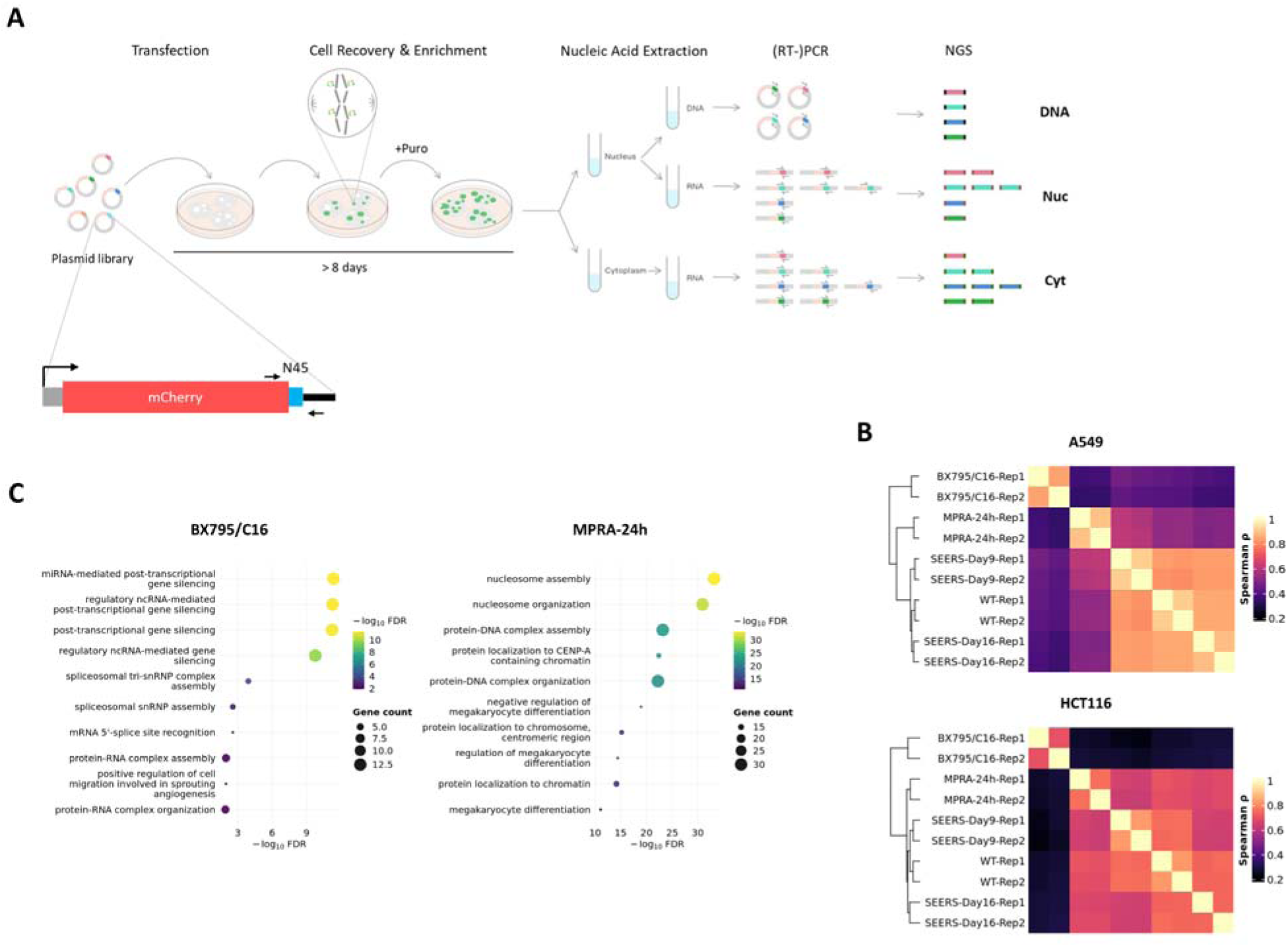
SEERS reduces cellular perturbations associated with standard MPRA workflows. (**A**) Schematic of the SEERS workflow and the mCherry reporter transcript. Small arrows indicate primer annealing sites used for amplicon sequencing of the N45 inserts. The SEERS vector is maintained episomally via the OriP/EBNA1 element and replicates once per cell cycle, enabling stable puromycin selection and enrichment of successfully transfected cells. (**B**) Transcriptome similarity matrix and heatmap computed using the 5,000 most variable genes across samples (by variance of log_2_[TPM + 1]), excluding rRNA and mitochondrial genes. WT denotes unmanipulated cells maintained under standard culture conditions. (**C**) Gene Ontology (GO) enrichment analysis (top 10 terms) of genes upregulated in the BX-795/C16-treated condition or in the standard MPRA condition (24-h collection), each compared with the WT control.

Consistent with this rationale, the transcriptomes of SEERS-selected cells were more similar to unperturbed wild-type (WT) controls than those obtained using either a conventional 24-h MPRA collection workflow or a workflow incorporating immunosuppressive small molecules (1 µM BX-795 and 1 µM C16) (**Fig. 1B**). Gene Ontology analysis of upregulated genes indicated that, although BX-795/C16 reduced immune-stress signatures, it concomitantly enriched pathways related to miRNA-mediated gene silencing and spliceosome assembly (**Fig. 1C**), changes that could confound functional interpretation of 3′UTR inserts. In contrast, the most significantly enriched terms in the 24-h MPRA workflow were dominated by nucleosome assembly (**Fig. 1C**), consistent with rapid chromatinization of plasmids upon nuclear entry^48^ and suggesting an additional workflow-associated source of confounding for assessing 3′UTR sequence function.

### Short 3′UTR sequence elements show widespread regulatory associations

We constructed an mCherry reporter driven by the EF-1α core promoter, in which a randomly synthesized 45-nt sequence (N45) was inserted immediately downstream of the mCherry stop codon (**Fig. 1A**). Analysis of the resulting library indicated that these random N45s provide sufficient sequence diversity to broadly encompass the short k-mer compositional patterns and nucleotide-level complexity observed in the proximal portions of native human 3′UTRs (**Supplemental Fig. S2**). Using this plasmid library (>10^6^ distinct inserts), we performed SEERS screens in both A549 and HCT116 cells. For each experiment, we collected nuclear DNA, nuclear RNA, and cytoplasmic RNA, and quantified the relative abundance of each N45 in each fraction by amplicon sequencing (DNA, Nuc, and Cyt) (**Fig. 1A**). Measurements were highly concordant across technical and biological replicates (**Supplemental Figs. S3 and S4**). Because A549 exhibited higher reproducibility across biological replicates, we first carried out downstream analyses using the A549 dataset.

To reduce contributions from sequencing artifacts and other technical noise, we removed N45s with extremely low DNA representation, yielding a dataset comprising ∼2 million N45s (**Supplemental Fig. S5**). We then normalized subcellular abundances by DNA representation and computed expression scores for each N45 in the nucleus and cytoplasm as Nuc score = log_2_(Nuc/DNA + 0.01) and Cyt score = log_2_(Cyt/DNA + 0.01). Comparing these scores revealed a global trend in which low-expression inserts tend to be nuclear enriched, whereas high-expression inserts tend to be cytoplasmic enriched (**Fig. 2A**). We further defined an overall Expression score = (Nuc score + Cyt score)/2 and an Export score = Cyt score − Nuc score.

**Fig. 2.**
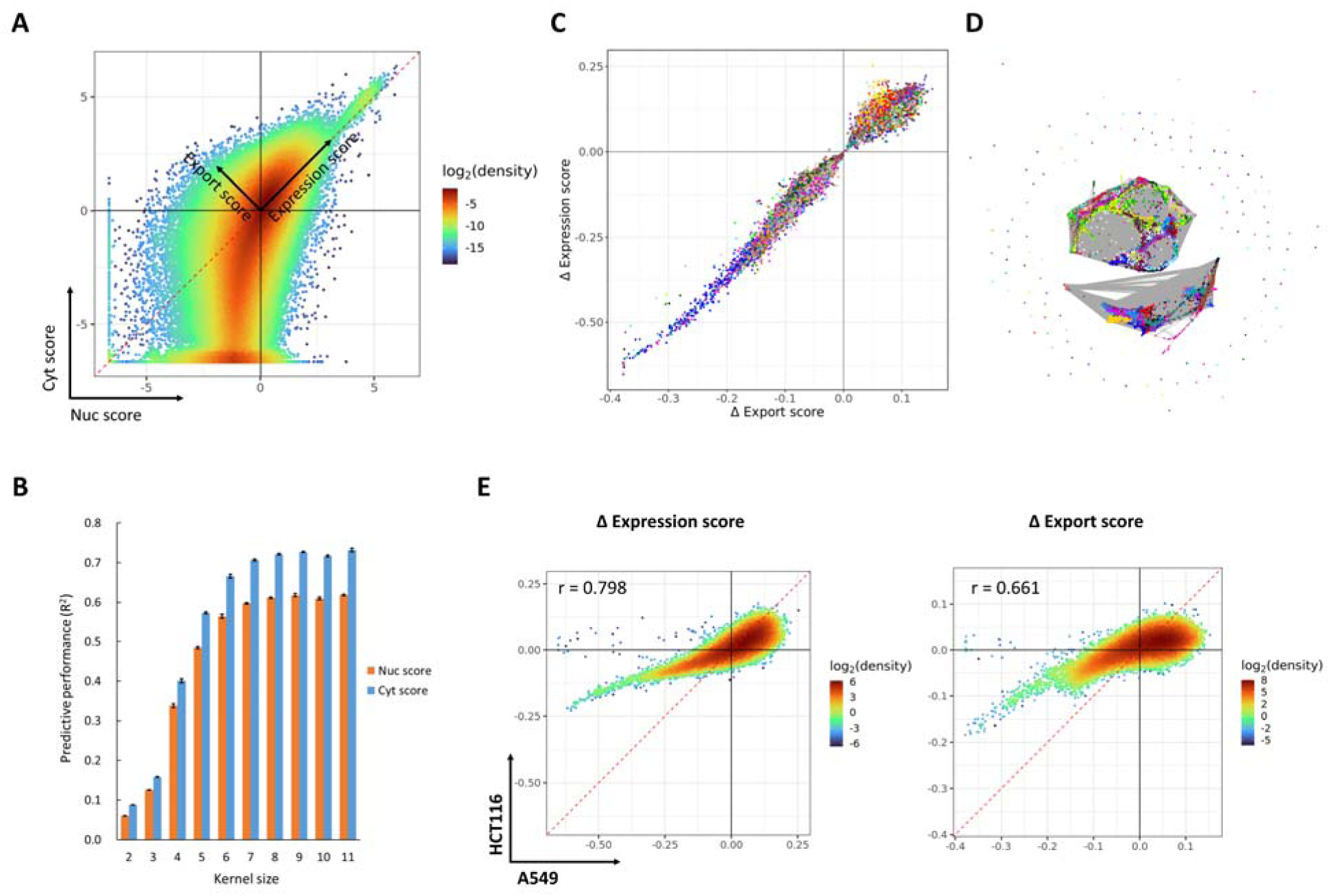
Short 3′UTR sequence elements show widespread regulatory associations. (**A**) Nuclear and cytoplasmic scores for the ∼2 million N45 inserts with the highest DNA abundance (proportion) in A549, computed as log_2_(Nuc/DNA + 0.01) and log_2_(Cyt/DNA + 0.01), respectively. (**B**) Effect of convolutional kernel size on the single-layer CNN “motif-counting” model. Each kernel size was evaluated in at least three independently trained models. Predictive performance is reported on the primary test set (n = 10,000) as Pearson r² (R²). (**C**) Distribution of regulatory properties for all 2∼8-mers showing significant regulatory correlations (*p* < 1 × 10; n = 36,945), colored by cluster assignment. (**D**) Clustering of the significant 2∼8-mers. Gray edge connect adjacent k-mers, defined as pairs differing by a single edit (edit distance = 1) with similar regulatory properties (0.5 < fold change < 2). k-mers were clustered from this adjacency graph using the Louvain algorithm. (**E**) Comparison of regulatory properties for all 8-mers (n = 65,536 between A549 and HCT116.

To more accurately benchmark model performance, we carried out an independent SEERS experiment in A549 using a small library of ∼3,000 N45 inserts but sequenced at substantially greater depth. This dataset showed high concordance between experimental replicates (r = 0.939 for Nuc score and r = 0.981 for Cyt score; **Supplemental Fig. S6A**). Importantly, sequences in this test set were well separated from the ∼2 million N45s used for training, with a minimum edit distance of 10 and a median edit distance of 15 (**Supplemental Fig. S6B**).

Using this stringent hold-out set, a simple, linear, single-layer CNN “motif-counting” model provided a good fit (best performance: Pearson R² = 0.619 for Nuc score and 0.732 for Cyt score), consistent with the view that much of the RNA regulatory code can be captured by the combined contributions of multiple short motifs.^42,49,50^ Notably, model performance plateaued when the convolution kernel length reached 8 nt (**Fig. 2B**), suggesting that the predominant, relatively independent sequence features (“regulatory elements”) are typically ≤8 nt, consistent with the characteristic lengths of miRNA seeds and many RNA-binding protein (RBP) binding motifs.^40,41,51^

Moreover, when the convolution kernel length was set to 2, 3, or 4 nt, predictive performance reached ∼12%, ∼22%, and ∼56%, respectively, of that achieved with an 8-nt kernel. This pattern indicates that even collections of very short sequence features on their own account for a substantial, quantifiable fraction of the regulatory “vocabulary” encoded in 3′UTRs, consistent with previous reports.^22,26^

Motivated by these observations, we systematically evaluated all 2∼8-nt k-mers for their associations with N45 regulatory phenotypes. For each k-mer, we compared the distributions of Expression score and Export score among N45s containing that k-mer with the corresponding distributions across all N45s, thereby obtaining a Δ Expression score and Δ Export score for that k-mer. This analysis revealed that ∼42% of all 2∼8-mers (36,945/87,376) exhibited significant regulatory associations (Mann-Whitney U test, p < 1 × 10^-6^; **Supplemental Table S1**). Using the Δ Expression score distribution of all 7-mers as a background, human miRNA seed 7-mers from TargetScan were significantly biased toward decreased expression (Mann-Whitney U test, p = 2.62 × 10^-9^; **Supplemental Fig. S7A**). In addition, N45s carrying k-mers corresponding to AU-rich elements (AREs), m6A-associated sites, or GU-rich elements showed significantly reduced Expression scores relative to background (Mann-Whitney U test, p < 2.05 × 10^-7^; **Supplemental Fig. S7B**). Together, these results indicate that SEERS robustly captures multiple well-established classes of human 3′UTR regulatory signals.^6,26^

Further integrating k-mer properties along the axes of expression level and subcellular localization revealed two broad functional groupings: (i) low-expression, nuclear-enriched elements and (ii) high-expression, cytoplasmic-enriched elements (**Fig. 2C**). This organization is consistent with a coupling between expression output and nuclear export/cytoplasmic enrichment in human cells.^35,52^

Finally, we performed Louvain clustering of significant 2∼8-mers using joint sequence and functional similarity, identifying 173 communities. Among these, 47 major communities contained 192∼2,200 k-mers each, whereas the remaining 126 communities were small (1∼5 k-mers). Despite this structure, extensive similarity between communities resulted in diffuse cluster boundaries (**Fig. 2D**). This pattern suggests that many significant k-mers may not correspond to unique, one-factor binding motifs, but instead reflect composite regulatory signals contributed by multiple RBPs and/or miRNAs, producing hybrid sequence signatures that approximate averaged binding preferences. Accordingly, we treated each k-mer as an independent regulatory element rather than collapsing them into larger motif patterns, as premature merging could introduce bias. Given the substantial motif similarity shared across many RBPs,^40,51,53^ we deliberately refrain from assigning individual regulatory elements to specific trans-acting factors or speculating on factor-specific mechanisms at this stage.

Notably, the 8-mer regulatory profiles were highly concordant between HCT116 and A549, with strong correlations for both Δ Expression score (r = 0.798) and Δ Export score (r = 0.661) (**Fig. 2E**). This cross-cell-line agreement suggests that a substantial fraction of the identified regulatory elements are shared between these epithelial cell lines. We therefore focus subsequent analyses on the A549 dataset and do not present a separate, parallel analysis for HCT116.

### Non-strand-specific 3**′**UTR elements are distinct from enhancer-like signals

Among all significant 2∼8-mers, ∼9% (3,208/36,945) lacked clear strand specificity, i.e., their regulatory properties were similar to those of their reverse-complement k-mers. Notably, ∼97% of these non-strand-specific k-mers (3,121/3,208) were positively associated with expression (**Fig. 3A**). To test whether these k-mers might instead report enhancer-like transcription factor (TF) signals, we repositioned the N45 insert from the 3′UTR to the 5′UTR of the mCherry reporter (immediately upstream of the start codon) (**Fig. 3B**), constructed a plasmid library containing ∼5 × 10^4^ N45 variants, and performed SEERS-5′UTR experiments in A549 cells. The SEERS-5′UTR measurements were highly reproducible across technical and biological replicates (**Supplemental Fig. S8**).

**Fig. 3.**
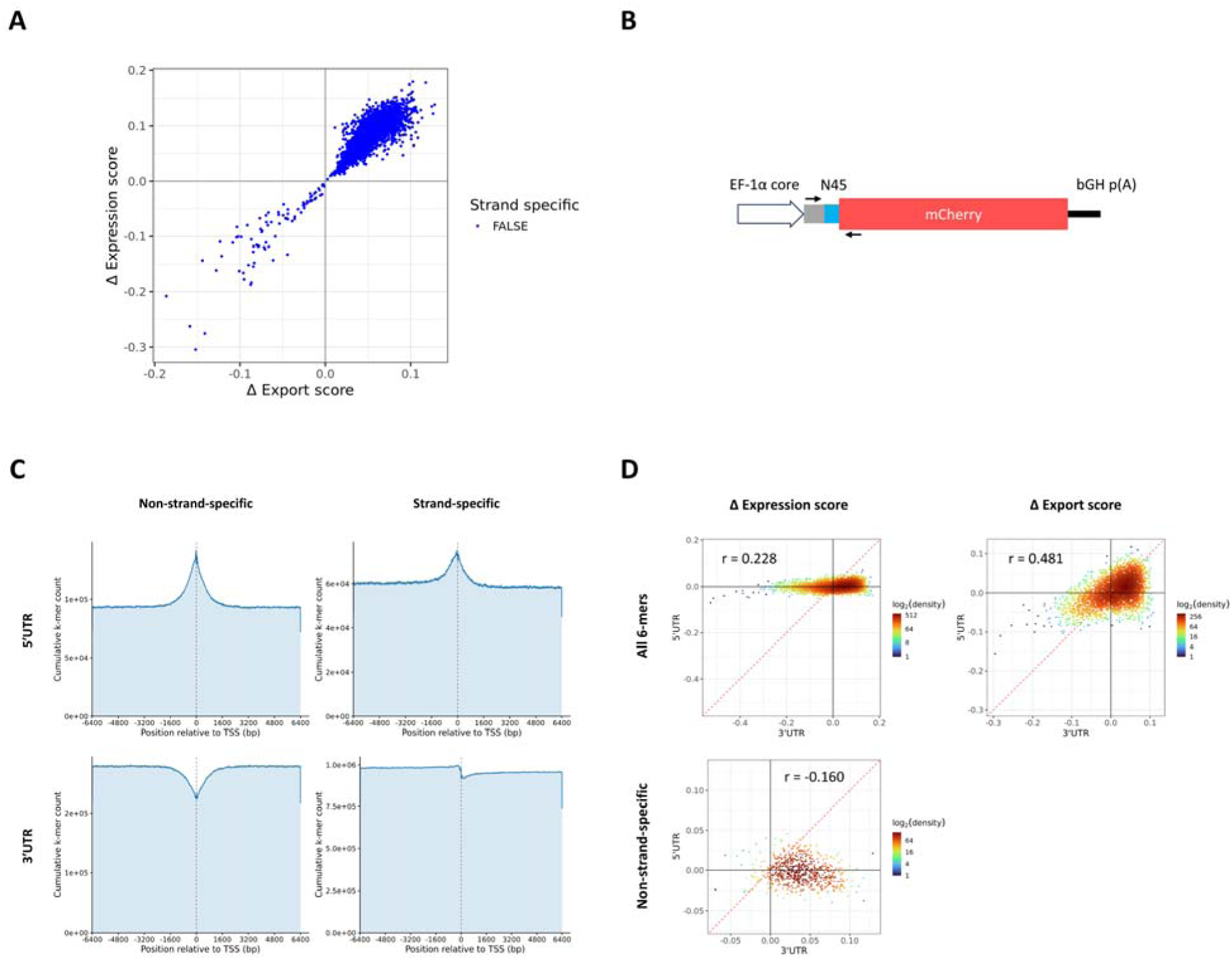
Non-strand-specific 3′UTR elements are distinct from enhancer-like signals. (**A**) Non-strand-specific subset (n = 3,208) among all 2∼8-mers showing significant regulatory correlations (p < 1 × 10^−6^; n = 36,945). For each k-mer and its reverse complement, the pair was classified as strand-specific if their regulatory properties differed substantially (fold change > 2 or < 0.5) and p < 0.01; otherwise, the pair was classified as non-strand-specific. (**B**) Schematic of th mCherry reporter configuration used in the SEERS-5′UTR experiment. (**C**) Enrichment of 6-mers with significant expression-enhancing effects (p < 0.01) in SEERS-3′UTR or SEERS-5′UTR experiments around GENCODE v49 annotated transcription start sites (TSSs) in the human genome. (**D**) Comparison of regulatory properties for all 6-mers (n = 4,096) between SEERS-3′UTR and SEERS-5′UTR experiments.

Because the library size was insufficient to support robust 7∼8-mer statistics, we analyzed 6-mers in the 5′UTR dataset. We identified 158 non-strand-specific and 124 strand-specific 6-mers that were significantly positively associated with expression (p < 0.01). These 6-mers were strongly enriched within ±1.6 kb of GENCODE v49 annotated transcription start sites (TSSs) in the human genome (**Fig. 3C**), consistent with enhancer/promoter-associated TF signals. In contrast, in the 3′UTR dataset, the 1,947 strand-specific 6-mers with significant positive associations showed no such enrichment, whereas the 493 non-strand-specific 6-mers were depleted near TSSs (**Fig. 3C**). Moreover, the regulatory properties of 6-mers in the 5′UTR and 3′UTR assays were not concordant (r = −0.160) (**Fig. 3D**). Together, these results argue that non-strand-specific elements detected in 3′UTRs do not primarily reflect enhancer-like transcriptional signals. Mechanistically, such orientation-insensitive behavior is compatible with models in which at least a subset of these elements are read out in the context of local RNA secondary structure or double-stranded RNA segments, rather than as canonical, directionally constrained single-stranded motifs.

Despite this distinction, when considering all 6-mers, the overall regulatory landscape showed modest similarity between 3′UTR and 5′UTR assays for both expression (r = 0.228) and localization (r = 0.481) (**Fig. 3D**), suggesting that 5′UTRs and 3′UTRs may share aspects of post-transcriptional regulation, particularly mechanisms related to nuclear-cytoplasmic partitioning.^2^

### U1 telescripting underlies strong repressive effects in 3**′**UTRs

In the k-mer analysis above, U1-like binding-site patterns (exemplified by AGGTAAGT and closely related k-mers) showed the strongest negative regulatory correlations, indicating that they represent one of the most prominent repressive and nuclear-enriching signals in 3′UTRs, consistent with prior reports that U1-associated sequences can profoundly shape transcript fate.^25,26,54,55^ However, a case study of an N45 insert harboring a U1 site and exhibiting one of the strongest negative effects on both Expression score and Export score revealed that its impact was not a simple, local reduction in RNA output. Instead, the data were most consistent with U1 snRNP-mediated telescripting: U1 binding near the 3′ end can suppress nearby premature cleavage and polyadenylation (PCPA), thereby promoting RNA polymerase II readthrough and shifting 3′-end formation to a more distal termination site.^56,57^

In this N45 example, RNA read coverage showed that the mCherry transcript did not terminate at its canonical poly(A)/termination site, but instead extended into the downstream EBNA1 region and ultimately terminated at the EBNA1 poly(A) site, yielding an abnormally long 3′UTR (**Fig. 4A**). In parallel, we detected a clear splice junction between the U1 site within the N45 and a downstream 3′ splice acceptor located 8 nt upstream of the EBNA1 start codon (sequence: TCTCTTTTAG/TGTGAATC[ATG]), indicating that, in this vector context, the U1 site can function as a cryptic 5′ splice site that pairs with a downstream acceptor to define an intron that is subsequently spliced out. Functionally, this configuration was accompanied by a marked reduction in mCherry expression.

**Fig. 4.**
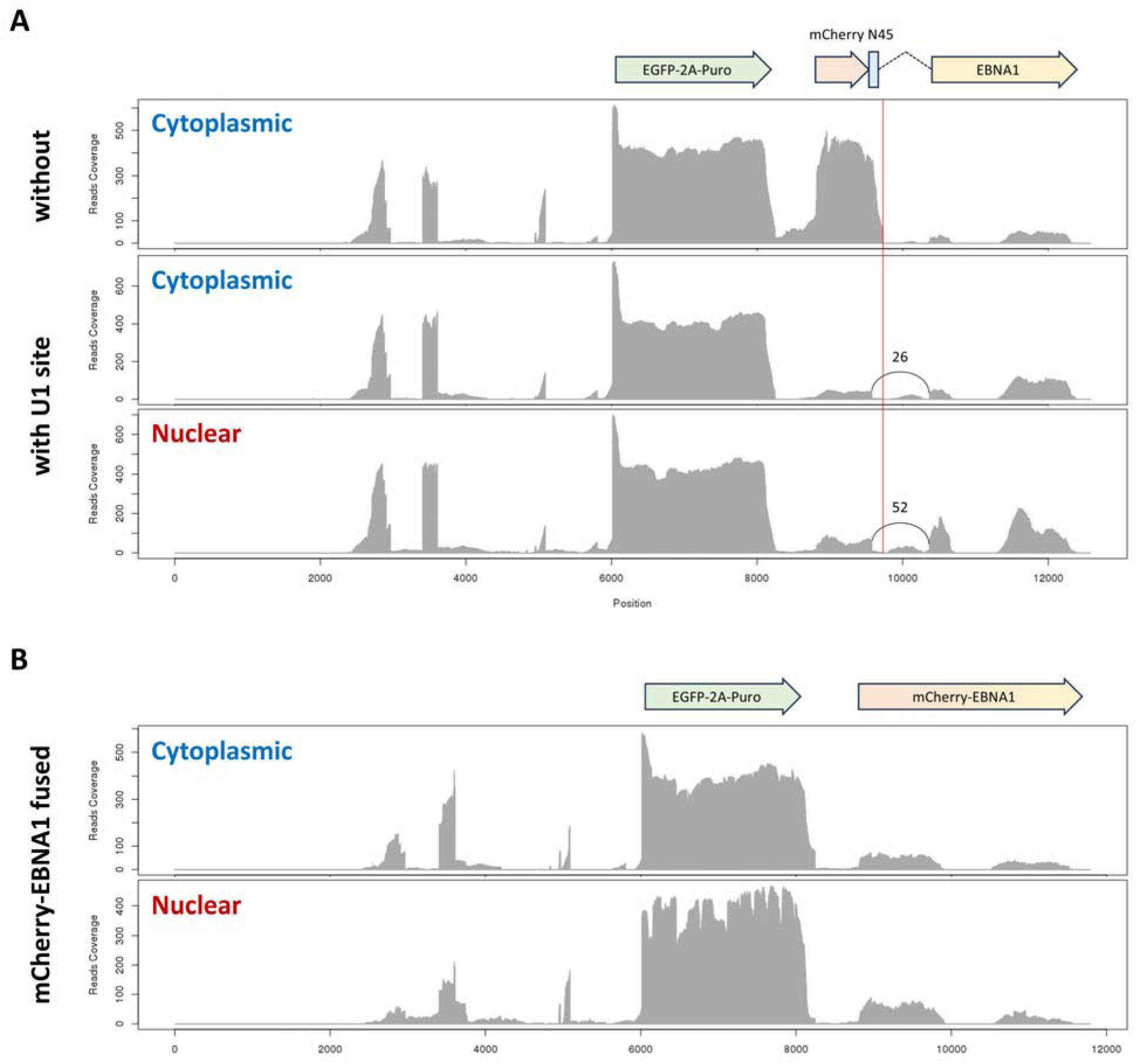
U1 telescripting underlies strong repressive effects in 3′UTRs. (**A**) RNA-seq read coverage across the SEERS reporter in constructs with or without a U1 site in the N45 insert. The vertical red line marks the transcription termination site that is suppressed by telescripting and traversed by elongating transcripts. Arcs denote splice junctions identified from RNA-seq; numbers indicate junction-supporting read counts. (**B**) RNA-seq read coverage after deletion of the splice-junction DNA sequence (i.e., fusion of mCherry and EBNA1), as in (A).

To distinguish whether the decrease in expression was driven primarily by the splicing event itself or by telescripting-associated readthrough and 3′-end repositioning, we deleted the DNA segment corresponding to the inferred intron (thereby eliminating this splicing path) and re-assayed mCherry expression under matched conditions. The deletion construct still produced a comparably strong reduction in mCherry expression (**Fig. 4B**), indicating that the dominant driver is not splicing per se, but rather U1-mediated readthrough leading to 3′UTR hyperextension and relocation of 3′-end formation. Taken together, these observations indicate that, in this reporter architecture, the strong negative effects associated with U1-related k-mers largely reflect telescripting-driven remodeling of transcript structure and 3′-end processing, rather than the more local, approximately additive behavior typical of many 3′UTR cis-elements. Given that U1-dependent telescripting is a well-established endogenous mechanism that can strongly influence transcript output by suppressing premature 3′-end formation,^56,57^ our results support the view that U1-related elements constitute a particularly potent class of regulatory signals in 3′UTRs, although their quantitative effects will depend on sequence context and the availability of downstream processing signals.

### Element effects are largely additive, with rare strong context-dependent modulation

To examine the extent to which the activities of regulatory elements depend on local sequence context, we trained an LSTM-based neural network, TALE (Transcript Abundance and Localization Estimator). Relative to the simple CNN “motif-counting” model, TALE improved predictive performance by ∼13% (increasing the Cyt score R^2^ from ∼0.73 to ∼0.86; **Fig. 5A**), indicating that contextual features and inter-element interactions make a measurable contribution to regulatory output. In addition, to benchmark TALE against more general MPRA modeling frameworks, we re-trained a DREAM challenge-style sequence model on our data using the same train/validation/test splits and evaluation protocol, and found that TALE consistently outperformed this baseline on both tasks (Pearson R^2^: 0.808 → 0.830 for Cyt score; 0.831 → 0.875 for Nuc score). Together, these comparisons suggest that while context-dependent effects are detectable and improve predictive performance over purely motif-counting and generic sequence models, their overall impact on 3′UTR-driven abundance and localization remains moderate.

**Fig. 5.**
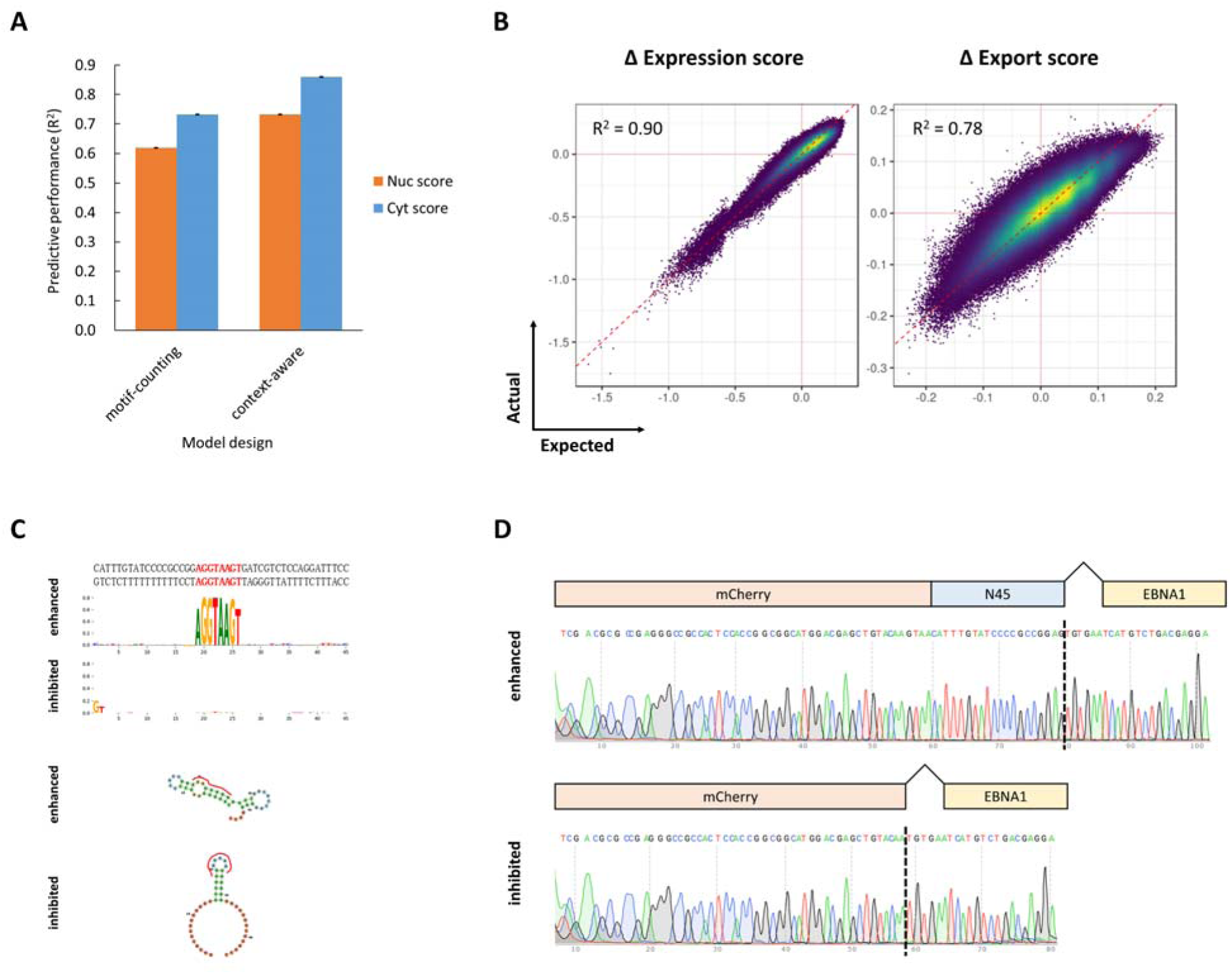
Element effects are largely additive, with rare strong context-dependent modulation. (**A**) Comparison of the best predictive performance of the LSTM-based context-aware model (TALE) and the CNN-based motif-counting model. Performance is reported as the coefficient of determination (R²) on the primary test set (n = 10,000). Each model was trained independently three times (n = 3). (**B**) Observed regulatory effects (Δ Expression or Export score) for any pairwise 5-mer combinations within N45 inserts (n = 524,800 pairs), compared with linear expectation values obtained by summing the corresponding single 5-mer effects. (**C**) Two N45 sequences generated by TALE-guided directed evolution that maximally enhance or maximally suppress the effect of the central U1 site. Nucleotide logo heights correspond to in silico saturated mutagenesis (ISM) scores. Predicted RNA secondary structures for both sequences were generated with RNAfold. (**D**) Experimental validation of the two TALE-designed N45s, including Sanger sequencing chromatograms of the corresponding cDNAs and schematic splice patterns. Vertical dashed lines indicate splice junction positions.

To test whether regulatory elements exhibit systematic cooperativity, we compared the joint effect of any two identical or different 5-mers within the same N45 to the linear expectation obtained by summing their individual effects (the dataset did not support a robust analogous analysis for 8-mers). Observed and expected effects were highly concordant (**Fig. 5B**), suggesting that most 5-mers act predominantly in an additive manner rather than through widespread strong synergy. Consistent with this, mapping the subset of 5-mer pairs whose joint effects exceeded additivity did not reveal an obvious clustering structure (**Supplemental Fig. S9**), implying that interactions among 3′UTR elements are generally weak and dispersed.

Because the U1-like site (AGGTAAGT) represents one of the strongest regulatory element classes, we used it as a case study to further probe how context modulates element usage. We fixed a U1 site at the center of N45 and, guided by TALE, performed in silico directed evolution on the remaining positions. The resulting “suppressive” context was predicted to fold into an RNA secondary structure effectively “locked” by a stem-loop, and the first two bases of N45 were evolved to GT (**Fig. 5C**), which together with the last two codons of mCherry (AAGTAA) created a new U1 site (AAGTAAGT) at the mCherry-N45 junction. This architecture was predicted to compete with the central U1 site for splice-site usage. Experimental validation of the evolved “enhancing” and “suppressive” N45s supported these predictions: in the enhancing context, the central U1 site was efficiently engaged as a 5′ splice site, whereas in the suppressive context, usage of the central U1 site was nearly undetectable and splicing was instead dominated by the newly created junctional U1 site (**Fig. 5D**). Taken together, these observations show that, in extreme sequence contexts, higher-order RNA structure and competition between neighboring functional sites can markedly enhance or attenuate the activity of individual regulatory elements.

### TALE prioritizes a subset of disease-associated 3**′**UTR variants

Based on our results in A549 and HCT116 cells, together with prior evidence that a substantial fraction of 3′UTR regulatory logic is shared across cell types from distinct tissue origins,^26^ we reasoned that TALE may be able to capture generalizable causal patterns underlying a subset of human 3′UTR variants.

To this end, we used TALE to predict the regulatory effects of 3′UTR single-nucleotide variants (SNVs) annotated in ClinVar. To maximize generalizability, for each SNV we evaluated its effect at every possible position within N45 and then took the median predicted effect across all positions as the final score. Using these scores, we found that the distributions of predicted effect sizes differed significantly between variants annotated as Pathogenic and Benign (Kolmogorov-Smirnov test, p = 0.000133). Taking the predicted effects of all 15,225 benign 3′UTR SNVs as an empirical null distribution, we found that 43 of 190 pathogenic 3′UTR SNVs exhibited significantly aberrant predicted effects (empirical p < 0.05) (**Fig. 6A**; **Supplemental Table S2**).

**Fig. 6.**
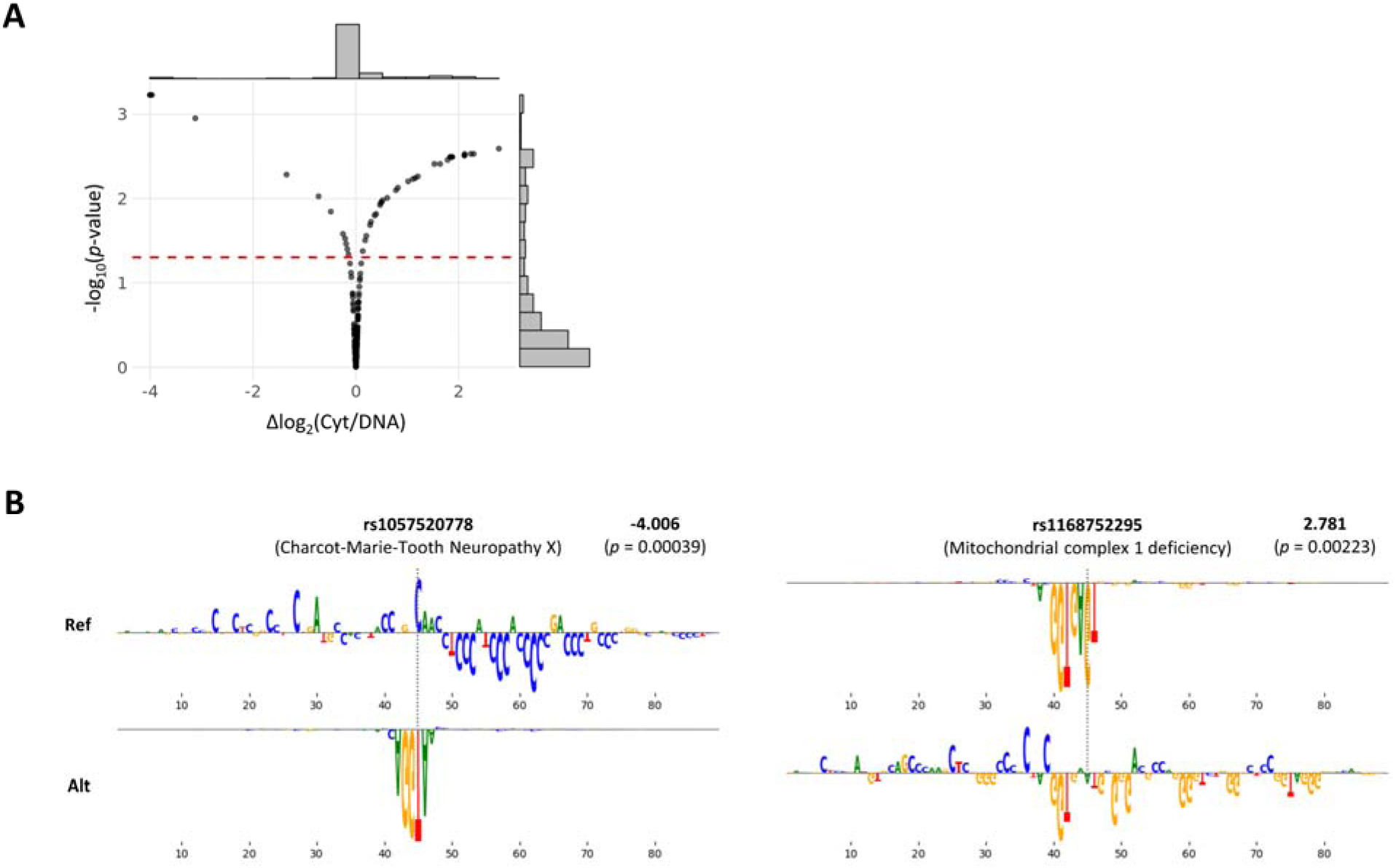
TALE prioritizes a subset of disease-associated 3′UTR variants. (**A**) Volcano plot of TALE-predicted mutation effects for all 190 pathogenic 3′UTR single-nucleotide variants (SNVs) in ClinVar. Mutation effect is defined as the difference in log2(Cyt/DNA) between the mutant and reference alleles (mutant − reference). The y axis shows −log10(empirical p-values), computed by comparing each pathogenic SNV’s predicted effect to a null distribution given by the predicted effects of all 15,225 benign 3′UTR SNVs in ClinVar. The red dashed line indicates the significance cutoff (p = 0.05) (**B**) The strongest TALE-predicted effects among pathogenic ClinVar 3′UTR SNVs are primarily driven by disruption or creation of 5′ splice sites. Nucleotide logo heights correspond to in silico saturated mutagenesis (ISM) scores. Upward logos indicate mutations predicted to decrease transcript abundance, whereas downward logos indicate increased abundance. Bold values denote TALE-predicted mutation effects (log2[Cyt/DNA]); parenthetical values report the corresponding empirical p-values derived from the benign-SNV null distribution described in (**A**).

Examination of the pathogenic SNVs with the strongest predicted effects revealed a degree of mechanistic convergence: a substantial fraction of these variants either create or disrupt 5′ splice sites (**Fig. 6B**), which could activate or suppress telescripting and thereby remodel 3′-end processing and overall mRNA architecture. This provides a concrete mechanistic basis for how subtle single-nucleotide changes within 3′UTRs can give rise to pronounced phenotypic consequences, consistent with previous reports.^26,58^ Notably, the enrichment of splice-site-altering variants also aligns with our broader inference that U1-related signals constitute a particularly potent class of regulatory elements in 3′UTRs.

## DISCUSSION

In this study, we introduce SEERS, an episome-based MPRA framework that enables large-scale measurement of 3′UTR sequence function while attenuating cellular perturbations associated with standard MPRA workflows. By coupling SEERS with a library of random 3′UTR inserts and context-aware modeling, we systematically mapped how short sequence features relate to two coupled phenotypic axes: RNA output and nuclear-cytoplasmic partitioning. Across ∼2 million synthetic inserts and two epithelial cell lines, we observed that a substantial fraction of short k-mers show reproducible regulatory associations, and that these associations largely organize along a dominant direction that links higher RNA abundance with cytoplasmic enrichment and lower abundance with nuclear enrichment. These results provide an empirical, sequence-level view of how 3′UTR information can be distributed across many weak contributors rather than a small number of dominant motifs.

A key implication of our measurements is that much of the regulatory behavior of these synthetic 3′UTRs can be approximated by additive composition of short elements. A simple “motif-counting” model captures a large portion of the variance in both abundance and localization, and direct additivity tests with 5-mers further support that most element pairs do not exhibit strong non-linear interactions. At the same time, the improved performance of a context-aware model (TALE), together with in silico directed evolution and experimental validation, indicates that context can matter in specific regimes: rare, extreme contexts can gate element usage through higher-order constraints such as RNA secondary structure or competition between nearby functional sites. Thus, the emerging picture is not that context is irrelevant. Rather, context dependence is typically modest and becomes prominent mainly when sequence arrangements create or suppress processing signals with large downstream consequences.

Our analysis highlights U1-related signals as an instructive example of how a short motif can dominate reporter output through changes in RNA processing. U1-like sites show the strongest negative associations in the 3′UTR assay, yet a detailed case study indicates that these effects frequently reflect U1 snRNP-mediated telescripting and associated remodeling of 3′-end formation and transcript architecture, rather than a purely local, approximately additive modulation of mRNA stability or export. This observation has two broader implications. First, it reinforces that 3′UTR “regulatory elements” can operate through qualitatively distinct mechanisms, including re-routing of cleavage/polyadenylation and splicing, whose quantitative effects may depend strongly on the availability of downstream processing signals. Second, it suggests that MPRA-based assays of 3′UTRs require particular care when sequences introduce cryptic splice sites or perturb 3′-end processing, because large apparent “element effects” may partly arise from transcript redefinition rather than modulation of the intended 3′UTR segment.^31^

SEERS complements and extends prior 3′UTR MPRA efforts that primarily interrogated naturally occurring sequences. Existing studies have successfully recovered canonical signals such as miRNA seed matches and AU-rich elements and have provided important insights into element function in physiological sequence contexts. However, reliance on endogenous sequence repertoires limits coverage of sequence space and makes it difficult to disentangle overlapping signals or to generalize across heterogeneous transcript architectures. In contrast, random libraries provide an approximately unbiased background against which many short signals can be evaluated systematically, and they enable data-hungry learning approaches to map sequence-to-phenotype relationships. Consistent with this, SEERS recapitulates multiple known classes of 3′UTR signals while also providing a global landscape of weak, distributed associations that are difficult to resolve from endogenous variation alone. Importantly, SEERS was designed to mitigate workflow-driven cellular responses, an increasingly recognized source of bias in MPRA experiments, and our transcriptome comparisons indicate that the selection-and-recovery strategy reduces perturbation relative to early collection or pharmacological suppression of immune signatures. Nevertheless, SEERS does not eliminate all potential confounders, and its episomal setting differs from the endogenous chromatin and transcriptional environment. These considerations should inform interpretation when extrapolating to native genes.

Several limitations and future directions follow from our design. First, episomal reporters decouple 3′UTR effects from endogenous promoter choice, chromatin state, and transcription kinetics, which is advantageous for isolating post-transcriptional contributions but may miss regulatory dependencies that arise from co-transcriptional coupling or isoform choice. Second, our primary readouts focus on nuclear-cytoplasmic partitioning and steady-state RNA abundance; additional assays that directly measure decay rates, translation, subcellular sub-compartment localization, or 3′-end usage would help connect sequence associations to specific mechanistic steps. Third, while we observe strong concordance between A549 and HCT116, broader surveys across additional cell types and perturbation conditions will be necessary to quantify which signals are broadly active versus context restricted.

Finally, TALE provides a practical bridge from synthetic sequence landscapes to human variation. Although trained in a cancer-derived cell line, the model prioritizes a subset of pathogenic ClinVar 3′UTR variants with unusually large predicted effects, many of which appear to act by creating or disrupting splice-site-like signals consistent with U1-dependent processing changes. This enrichment suggests that a portion of disease-relevant 3′UTR variation may operate through discrete processing switches rather than subtle tuning of stability, and it motivates targeted experimental validation in endogenous contexts. More generally, integrating high-throughput synthetic measurements with context-aware prediction offers a testable framework for interpreting noncoding variation in 3′UTRs and for designing sequences with desired expression and localization properties.

## METHODS

### SEERS-3′UTR vector and library construction

The SEERS-3′UTR vector (LN110; also referred to as SEERS3p) was derived from pCXLE-EGFP (Addgene #27082). Briefly, the EGFP open reading frame was extended by appending a P2A-PuroR sequence, and an mCherry expression cassette was subsequently inserted downstream of the EGFP-P2A-PuroR cassette (see **Supplemental Table S3** for sequences of all plasmids and oligonucleotides).

N45 double-stranded DNA insertion fragments were generated by a single-cycle PCR using oligonucleotides 110_HindIII_N45 and bGH_R2 (0.5 µM each) with NEBNext Ultra II Q5 Master Mix (NEB) under the following conditions: 98°C for 30 s, 66°C for 2 min, and 72°C for 2 min. The resulting unpurified PCR products were assembled into LN110 at a HindIII site adjacent to the mCherry stop codon by Gibson assembly. Correct insertion was confirmed by Sanger sequencing (10/10 clones). Unless otherwise indicated, Gibson assemblies were performed using NEBuilder HiFi DNA Assembly Master Mix (NEB) or equivalent reagents. Assembled plasmids were transformed into XL10-Gold competent cells, plated on LB agar containing carbenicillin (100 µg/mL), and incubated at 37°C overnight. Plasmids were prepared using an EndoFree Maxi Plasmid Kit (Tiangen).

### Cell culture and transfection

A549 and HCT116 cell lines (obtained from the Cell Resource Center, Peking Union Medical College, and BNCC, respectively) were authenticated by STR profiling and confirmed to be mycoplasma-free. Cells were maintained in DMEM/F-12 (Procell) supplemented with 10% fetal bovine serum (Procell) and 1× penicillin–streptomycin–amphotericin B solution (Solarbio). SEERS libraries were co-transfected with pCXWB-EBNA1 (Addgene #37624) at a 4:1 molar ratio. Transfections were performed using Lipofectamine 3000 (Invitrogen) or TransIT-X2 (Mirus). To minimize cytotoxicity, transfection reactions (Opti-MEM, DNA, and transfection reagents) were typically scaled to 50% of the manufacturer’s recommended amounts. 24∼48 hours post-transfection, puromycin (Beyotime; 1 µg/mL) was added to the culture medium, and selection was continued until day 9∼11 post-transfection.

For SEERS-3′UTR experiments comprising ∼2 million N45 variants (L5), A549 experiments were performed in two batches: batch 1 (T1) included four biological replicates (two 10-cm dishes per replicate), and batch 2 (T2) included two biological replicates (LN and XH, seven 10-cm dishes per replicate). HCT116 experiments were performed in three batches: batch 1 (T2) included two biological replicates (one 10-cm dish per replicate), batch 2 (T3) included one biological replicate (two 10-cm dishes), and batch 3 (T5) included one biological replicate (four 10-cm dishes). For SEERS experiments comprising ∼3,000 N45 variants (L6), transfections were performed in a single 10-cm dish.

### Nuclear/cytoplasmic fractionation and nucleic acid extraction

Cells were dissociated with TrypLE Express (Gibco), washed with DPBS, and pelleted at 300 ×g for 2 min. For every ∼1×10^7^ cells, pellets were resuspended in 200 µL ice-cold RLN2 buffer (DPBS, 0.5% v/v Nonidet P-40) and incubated for 5 min. Lysates were gently pipetted and centrifuged at 500 ×g for 2 min. Approximately 180 µL of the supernatant was collected as the cytoplasmic fraction and mixed with 400 µL RLT Plus buffer (Qiagen). The nuclear pellet was washed once with 1 mL RLN2 buffer and once with 1 mL DPBS, then resuspended in 600 µL RLT Plus buffer and homogenized by repeated pipetting. DNA and RNA were co-purified from cytoplasmic and nuclear fractions using the AllPrep DNA/RNA Mini Kit (Qiagen) or AllPure Cell Kit (Magen). Fraction purity was further assessed by RNA-seq, using mitochondrial transcripts and snoRNAs as markers for cytoplasmic and nuclear fractions, respectively.

### Standard RNA-seq and alignment

All RNA-seq library preparation and sequencing were performed by Novogene. Where indicated, libraries were strand-specific. Sequencing was carried out on an Illumina NovaSeq platform using 2× 150 bp paired-end reads. Raw reads were quality-filtered and adapter-trimmed with fastp, then aligned to the human reference genome (hg38; chromosomes only) using STAR^59^. Gene-level read counts were obtained with Subread^60^ using GENCODE v49 basic annotations. Reads overlapping multiple meta-features (genes) were assigned to the gene with the largest overlap, or split evenly among genes with equal overlap. Because rRNA depletion during library preparation is not complete, all ribosomal RNA genes and related pseudogenes were excluded when calculating gene expression (TPM) to minimize systematic distortion of expression estimates for other genes. For similar reasons, all mitochondrial genes were also excluded.

### GO enrichment analysis

Upregulated genes in the target samples were defined as genes with TPM fold change > 16 relative to control samples and an absolute TPM > 8 in the target samples. This gene set was subjected to Gene Ontology (GO) enrichment analysis using clusterProfiler^61^ in R, with org.Hs.eg.db as the annotation database. Analyses were performed using ENSEMBL gene identifiers and restricted to the Biological Process (BP) ontology. Multiple testing correction was carried out using the Benjamini-Hochberg method. For the enrichment test, the background gene universe was defined as all genes detected in either the target or control samples (i.e., the union of expressed genes across the two conditions), using ENSEMBL identifiers. No additional p-value or q-value thresholds were applied at the enrichment step (i.e., all GO terms were retained for downstream filtering/visualization).

### Cell perturbation and toxicity comparison assay

Experiments were performed in both A549 and HCT116 cells with two biological replicates per cell line. Cells were transfected with the SEERS3p-L4 plasmid library (∼1×10^5^ N45 variants) using Lipofectamine 2000 (Thermo Fisher Scientific) according to the manufacturer’s instructions, without co-transfection of pCXWB-EBNA1. At 24 h post-transfection, one set of cells was harvested as a control for the standard MPRA workflow. In parallel, for the small-molecule inhibitor control, culture medium was replaced with medium containing BX-795 (1 µM) and C16 (1 µM), and cells were harvested after an additional 24 h. For the SEERS workflow condition, culture medium was replaced with medium containing puromycin (1 µg/mL) at 24∼48 h post-transfection, and cells were maintained under continuous selection and harvested at day 9∼16 post-transfection. Unmanipulated wild-type (WT) cells were collected in parallel at the corresponding time points.

Total RNA was extracted using the AllPure Cell Kit (Magen) and subjected to standard RNA-seq at Novogene. Following standard RNA-seq processing as described above, the 5,000 most variable genes across samples (by variance of log2[TPM+1]) were selected, pairwise expression-profile similarity (spearman, ρ) was computed, and the resulting similarity matrix was visualized as the heatmap shown in the manuscript.

### SEERS-3′UTR amplicon sequencing

We used two amplicon-sequencing workflows (Methods 1 and 2), with Method 2 representing an updated protocol. Datasets generated from August 2025 onward were processed using Method 2. Comparative analyses indicated that the two workflows did not introduce systematic differences overall; however, Method 2 mitigated an artifactual enrichment of poly(A) and A-rich elements in nuclear/cytoplasmic distributions observed in Method 1 (**Supplemental Fig. S10**), potentially arising from strand invasion of the oligo(dT) reverse-transcription primer at internal poly(A) tracts within the N45.

Method 1. Total RNA was reverse transcribed into cDNA using HiScript III RT SuperMix for qPCR (Vazyme) according to the manufacturer’s instructions. The N45 region was amplified from 1 µg genomic DNA or 5 µL cDNA in a 50 µL PCR reaction using NEBNext Ultra II Q5 Master Mix (NEB) and primers P5_N8_RD1_110_F and P7_N8_RD2_110_R (1 µM each). Cycling conditions were 98°C for 30 s; 25∼30 cycles of 98°C for 10 s, 65°C for 30 s, and 68°C for 45 s; followed by a final extension at 65°C for 5 min. Amplicons of the expected size (295 bp) were gel-excised and purified using the HiPure Gel Pure Micro Kit (Magen).

Method 2. Total RNA was reverse transcribed using Maxima H Minus Reverse Transcriptase (Thermo Scientific) with 1 µM gene-specific primer (SEERS3p_GSP), according to the manufacturer’s instructions. The N45 region was amplified from 1 µg genomic DNA or 10 µL cDNA in a 50 µL reaction using KAPA HiFi HotStart ReadyMix (Roche) and primers NR1_SEERS3p_F and NR2_SEERS3p_R (1 µM each). Cycling conditions were 95°C for 3 min; 21 cycles of 98°C for 20 s and 65°C for 45 s; followed by a final extension at 65°C for 5 min. For indexing, 1 µL of the unpurified first-round PCR product was re-amplified in a 50 µL reaction with Illumina Nextera index primers (e.g., N522 and N729) under the following conditions: 95°C for 3 min; 8 cycles of 98°C for 20 s and 65°C for 45 s; followed by a final extension at 65°C for 5 min. Final PCR products were purified directly using the HiPure Gel Pure DNA Mini Kit (Magen).

Purified libraries were sequenced by Novogene on an Illumina NovaSeq platform using 2× 150 bp paired-end reads. For each biological sample and fraction, multiple independent subsamplings and library preparations/sequencing runs were performed as technical replicates until replicate concordance plateaued, indicating saturation with respect to sampling and sequencing depth.

### SEERS data processing

Paired-end reads were first merged into single reads using NGmerge^62^. Adapter sequences flanking the N45 insert were then removed using the Biostrings R package. N45 inserts were counted and converted to within-sample proportions, yielding an “N45 proportion file” for each sample.

For libraries sequenced across multiple runs, N45 proportions were combined using run-specific sequencing depth as weights. For the same biological sample processed in multiple library-preparation batches, N45 proportions were combined using the corresponding nucleic-acid input amounts as weights. For the same plasmid library profiled across multiple biological samples within a given cell line, N45 proportions were combined using the number of culture dishes as weights.

N45s falling below the inflection point of the ranked DNA-abundance distribution (i.e., the “cliff” in **Supplemental Fig. S5**) were discarded, and the remaining N45 proportions were renormalized. Scores were then computed as follows: Cyt score = log2(Cyt/DNA + 0.01), Nuc score = log2(Nuc/DNA + 0.01), Expression score = (Cyt score + Nuc score)/2, and Export score = Cyt score − Nuc score, where Cyt and Nuc denote the N45 proportions in cytoplasmic and nuclear RNA samples, respectively, and DNA denotes the N45 proportion in the cellular DNA sample.

### SEERS k-mer analysis and clustering

For each k-mer, we first identified all N45 sequences in the SEERS dataset containing that k-mer. These N45s were then compared to the full N45 population using a Mann-Whitney U test to evaluate differences in the median Expression score and Export score, yielding Δ Expression score, Δ Export score, and corresponding p-values.

For a pair of reverse-complementary k-mers, if the two associated N45 subsets differed significantly (p < 0.01) and the effect size fell outside a 0.5∼2 fold-change window, the pair was classified as strand-specific.

For clustering, two k-mers were defined as adjacent if they differed by a single edit (edit distance = 1) and exhibited similar regulatory properties, with both Δ Expression score and Δ Export score within a 0.5∼2 fold-change window. An adjacency matrix was constructed accordingly, and all 2∼8-mers were clustered in R using igraph^63^ with cluster_louvain (resolution = 4). Network layouts were generated using layout_with_drl for visualization.

### Construction and training of neural networks

We merged the A549 and HCT116 SEERS datasets and retained only N45 inserts that were measured in both cell lines. Although most inserts were 45 nt in length (91.8%), insert lengths ranged from 6 to 128 nt. Because inserts longer than 46 nt were rare (∼0.7%), we standardized all sequences to a fixed length of 46 nt by truncating inserts >46 nt and padding shorter inserts with zeros at the 3′ end. Unless otherwise noted, these fixed-length inputs are referred to as N45.

From the resulting N45 set, we randomly held out 10% of sequences as a validation set and used the remaining sequences for training. An independent, deeply sequenced small-scale SEERS experiment in A549 (L6; ∼3,000 N45s; described above) was used as an independent test set, with no overlap among the training, validation, and test splits.

All models were implemented and trained in PyTorch. Training used the largest batch size permitted by the available GPU memory (18 GB; typically 2^14^). Models were optimized by minimizing mean absolute error (MAE) on the validation set, with early stopping to prevent overfitting. We used the Adam optimizer with a OneCycle learning-rate schedule, with the maximum learning rate set to 2^−9^ and the initial learning rate determined by max_lr/div_factor. This configuration yielded stable training dynamics and robust performance.

The best-performing architecture was an LSTM-based context-aware model, TALE. Inputs were first represented in one-hot format and converted to integer token IDs using the first four channels (A/C/G/T). Positions that were all-zero across these four channels, covering ambiguous bases and padding, were mapped to a single “N” token. An embedding layer then mapped the tokenized 46-nt sequence to a 46 × 5 representation. The embedded sequence was passed through two unidirectional LSTM layers with 128 and 64 hidden units, respectively, both returning hidden states at each time step, followed by dropout (rate = 0.5). The sequence of hidden states was flattened and fed into a fully connected layer with 128 units (ReLU activation) followed by dropout (rate = 0.5). A linear output layer jointly predicted four phenotypes: Nuc score and Cyt score in A549 and HCT116 (output order followed the dataset definition).

As a baseline “motif-counting” model, we implemented a CNN consisting of the same embedding layer, a 1D convolutional layer (stride = 1, ReLU activation), a global average pooling layer, and a linear output layer. For each convolutional kernel size, the number of filters and the learning rate were tuned independently.

### SEERS-5′UTR experiment

To enable cloning of 5′UTR inserts upstream of the reporter, the LN110 backbone was modified by restriction digestion and Gibson assembly to replace the AgeI-PstI fragment, introducing an XhoI site immediately 5′ of the mCherry start codon, yielding the SEERS5p vector. The 5′UTR insert was generated by a single-cycle PCR using KAPA HiFi HotStart ReadyMix with 1 µM each of oligonucleotides 5p_N45_F and 5p_N45_R (98°C for 40 s, 69°C for 30 s, 65°C for 5 min). The resulting unpurified double-stranded DNA product was inserted into the XhoI site of SEERS5p by Gibson assembly. Insertion efficiency was assessed by Sanger sequencing (10/10 clones verified).

Approximately 350 ng of purified Gibson assembly product was transformed into 800 µL XL10-Gold competent cells and plated across sixteen 10-cm LB-agar plates supplemented with carbenicillin for overnight growth. The following day, the plasmid library was isolated using the TIANGEN EndoFre Mini Plasmid Kit II.

A549 cells were co-transfected in two 10-cm dishes with the plasmid library and pCXWB-EBNA1 at a 10:1 molar ratio using TransIT-X2 (Mirus). At 24 h post-transfection, puromycin was added to 1 µg/mL. Cells were maintained under continuous selection and harvested on day 9 post-transfection. Nuclear/cytoplasmic fractionation and DNA/RNA extraction were performed as in the SEERS-3′UTR experiment. NGS library preparation was likewise performed as for the SEERS-3′UTR, except that SEERS3p_GSP was replaced with SEERS5p_GSP and primer pair NR1_SEERS3p_F/NR2_SEERS3p_R was replaced with NR_SEERS5p_F/NR_SEERS5p_R.

### In silico mutagenesis and directed evolution

To compute in silico saturated mutagenesis (ISM) scores for each nucleotide in an N45 sequence, we mutated the nucleotide to each of the other three possible bases, generating three single-mutant N45s. TALE was then used to predict the change in regulatory phenotype (Expression score or Export score) between each mutant and the original sequence (mutant − reference). The median of these three predicted differences was taken as the ISM score for that nucleotide.

For directed evolution, we enumerated all possible single-nucleotide saturation point mutants of a given starting N45 sequence and predicted their Expression score or Export score values with TALE. The mutant closest to the desired evolutionary target (e.g., maximal Expression score) was selected as the input for the next round of mutagenesis. This iterative procedure was repeated until no further improvement in sequence fitness could be achieved.

### Validation of U1-binding site context

To generate 3′UTR inserts containing either an enhanced or inhibited U1-binding site context, double-stranded DNA inserts were produced by a single-cycle PCR using KAPA HiFi HotStart ReadyMix with 1 µM each of oligonucleotides U1_enhanced (or U1_inhibited) and H1_R (95°C for 3 min; 98°C for 20 s; 65°C for 15 min; hold at 15°C). The unpurified PCR product was inserted into the HindIII site of the LN110 vector by Gibson assembly. Multiple clones were screened and verified by Sanger sequencing and Nanopore sequencing, and endotoxin-free plasmids (SEERS3p-U1e and SEERS3p-U1i) were prepared using the TIANGEN EndoFre Mini Plasmid Kit II.

A549 cells in 6-well plates were transfected with SEERS3p-U1e or SEERS3p-U1i (one well per construct) using TransIT-X2 (Mirus). At 24 h post-transfection, puromycin was added to 1 µg/mL, and cells were maintained under continuous selection until harvest on day 13 post-transfection. Cells were lysed directly in the wells, and genomic DNA and total RNA were isolated using the AllPure Cell Kit (Magen) according to the manufacturer’s instructions.

For cDNA synthesis, 2.19 µg RNA per sample was reverse-transcribed using the HiScript III 1st Strand cDNA Synthesis Kit (Vazyme) with the gene-specific primer EBNA1_5p_R1. The resulting cDNA was diluted 1:10, and 10 µL was amplified by PCR using KAPA HiFi HotStart ReadyMix with 1 µM primers mCherry_qF and EBNA1_5p_R2 (95°C for 3 min; 30 cycles of 98°C for 20 s and 65°C for 30 s; 65°C for 5 min; hold at 15°C). PCR products were subsequently subjected to Sanger sequencing using primer pDsRED-express-C1-F.

### Assessment of human 3′UTR variant effects

For each hg38 SNV, we extracted 44 nucleotides upstream and 44 nucleotides downstream of the variant to generate 89-nt reference (Ref) and alternative (Alt) allele sequences. Each 89-nt sequence was then tiled into 45 overlapping 45-mers using a 1-nt step size. TALE was used to predict the regulatory phenotype for each 45-mer. Ref and Alt 45-mers at matching window positions were compared pairwise to obtain Δ Cyt/DNA for each window. The median of these window-level Δ Cyt/DNA values was reported as the final estimate of the SNV effect.

### Data access

All raw sequencing data generated in this study have been deposited in the Genome Sequence Archive for Human (GSA-Human; https://ngdc.cncb.ac.cn/gsa-human/) under accession number HRA008408. All processed data and code are available at https://github.com/gao-lab/SEERS.

## Supporting information

Supplemental Infomation

## Acknowledgments

We thank Drs. C. Li and L. Kong, as well as their labs for their generous supports in experimental facilities. We thank X.W. Liu and L.M. Xiao for their artistic inspirations. This work was supported by funds from National Key Research and Development Program (grant no. 2016YFC0901603) and State Key Laboratory of Gene Function and Modulation Research (formerly State Key Laboratory of Protein and Plant Gene Research). Part of the analyses was supported by the Computing Platform of the Center for Life Sciences of Peking University and the High-performance Computing Platform of Peking University.

## Author contributions

N.L., G.G., and L.C.R. conceived the study. N.L. developed SEERS and conducted the experiments. N.L., J.Y.L., and Y.D. analyzed the experimental data. J.Y.L. and N.L. constructed the neural network models and performed downstream analyses. N.L. wrote the manuscript. G.G., L.C.R., J.C., and Y.W. revised the manuscript.

