## Supplemental Infomation for "A systematic delineation of 3′UTR regulatory elements and their contextual associations"

Nan Liang *et al.*

**This PDF file includes:**

Supplemental Figs. S1 to S10

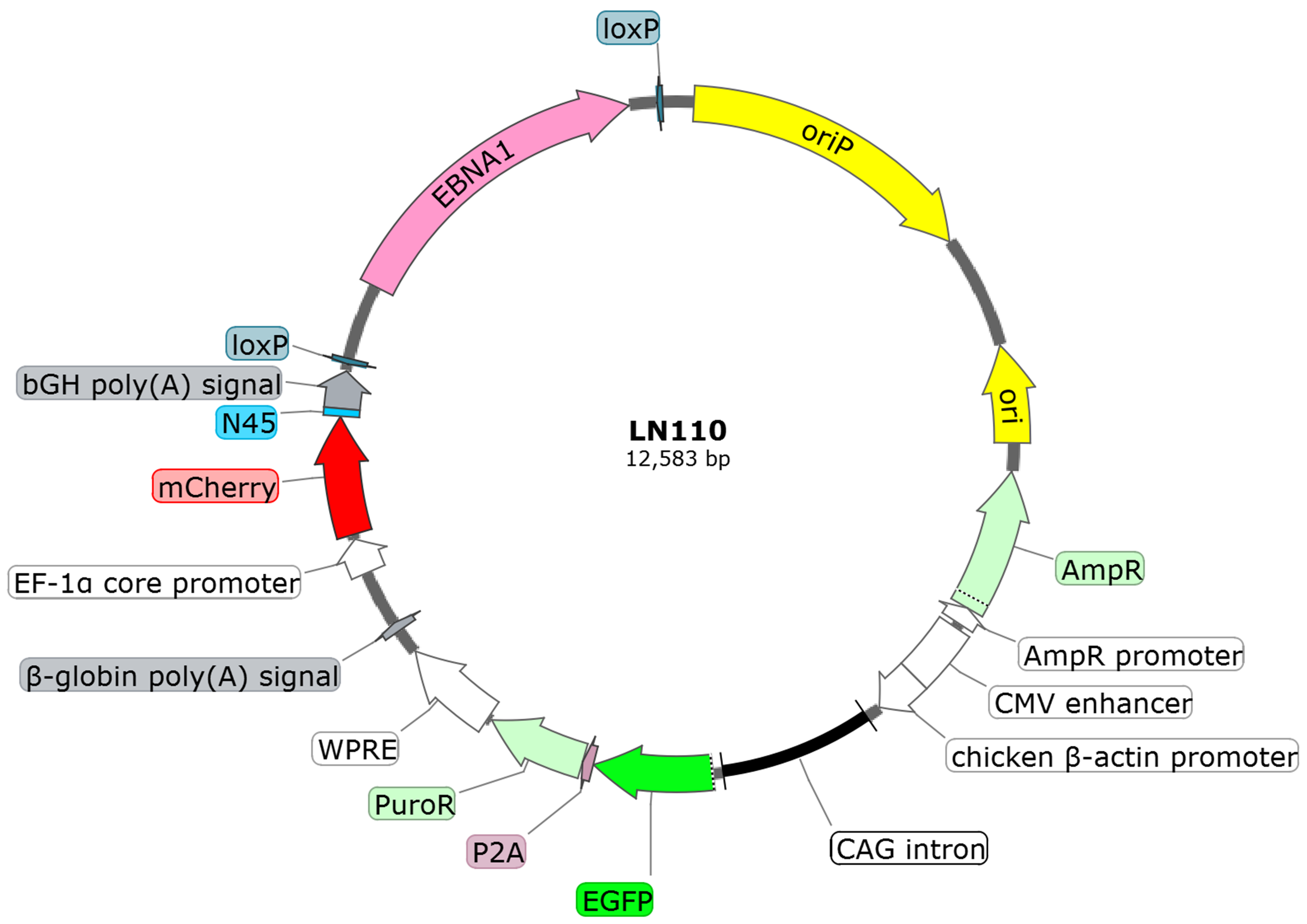

**Supplemental Fig. S1: Map of the SEERS3p (LN110) vector with the N45 insert.** Schematic of the SEERS3p (LN110) episomal reporter plasmid and its major functional modules. The vector carries the EBNA1/oriP elements to enable episomal maintenance through chromosome tethering and replication/segregation with the cell cycle. A CAG-driven EGFP-P2A-PuroR cassette allows visualization and puromycin selection of transfected cells, with transcript stabilization/enhancement conferred by WPRE and a β-globin poly(A) signal. The mCherry reporter is transcribed from a human EF-1α core promoter and terminated by a bGH poly(A) signal. The randomly synthesized 45-nt insert (N45) is positioned immediately downstream of the mCherry stop codon to serve as the 3′UTR insertion for SEERS screening and amplicon-based quantification.

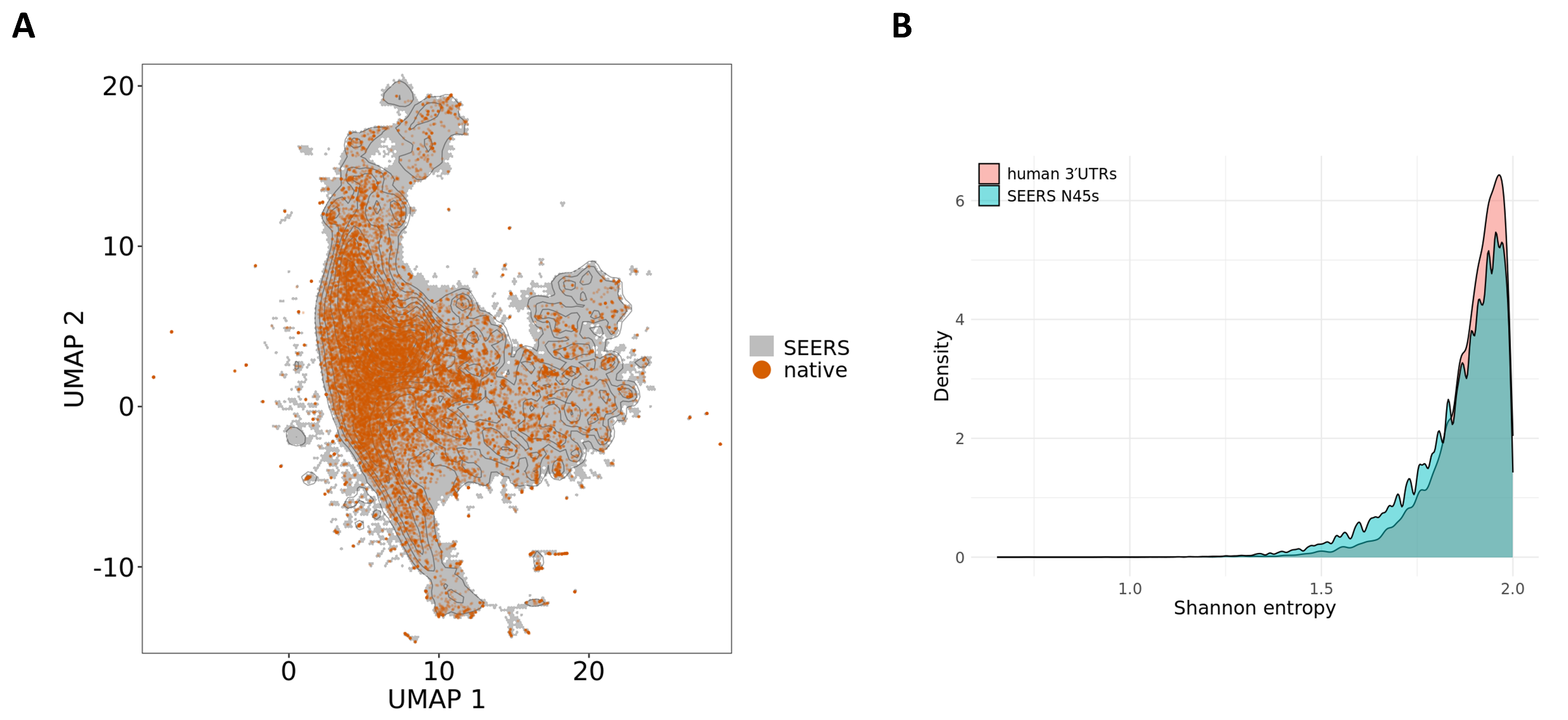

**Supplemental Fig. S2: Synthesized random N45 sequences show high complexity and broad coverage of 3′UTR sequence space.** (**A**) Comparison of sequence-space distributions between the randomly synthesized N45 library used in SEERS (background; gray) and annotated human 3′UTR sequences from GENCODE v49 (first 45 nt of each 3′UTR; orange points). Sequences were converted to 8-mer frequency vectors (L1-normalized) and visualized by UMAP dimensionality reduction. Density contours indicate the distribution of the N45 background. (**B**) Shannon entropy distributions of random N45s and native human 3′UTR sequences (first 45 nt), providing an information-theoretic measure of sequence diversity and complexity.

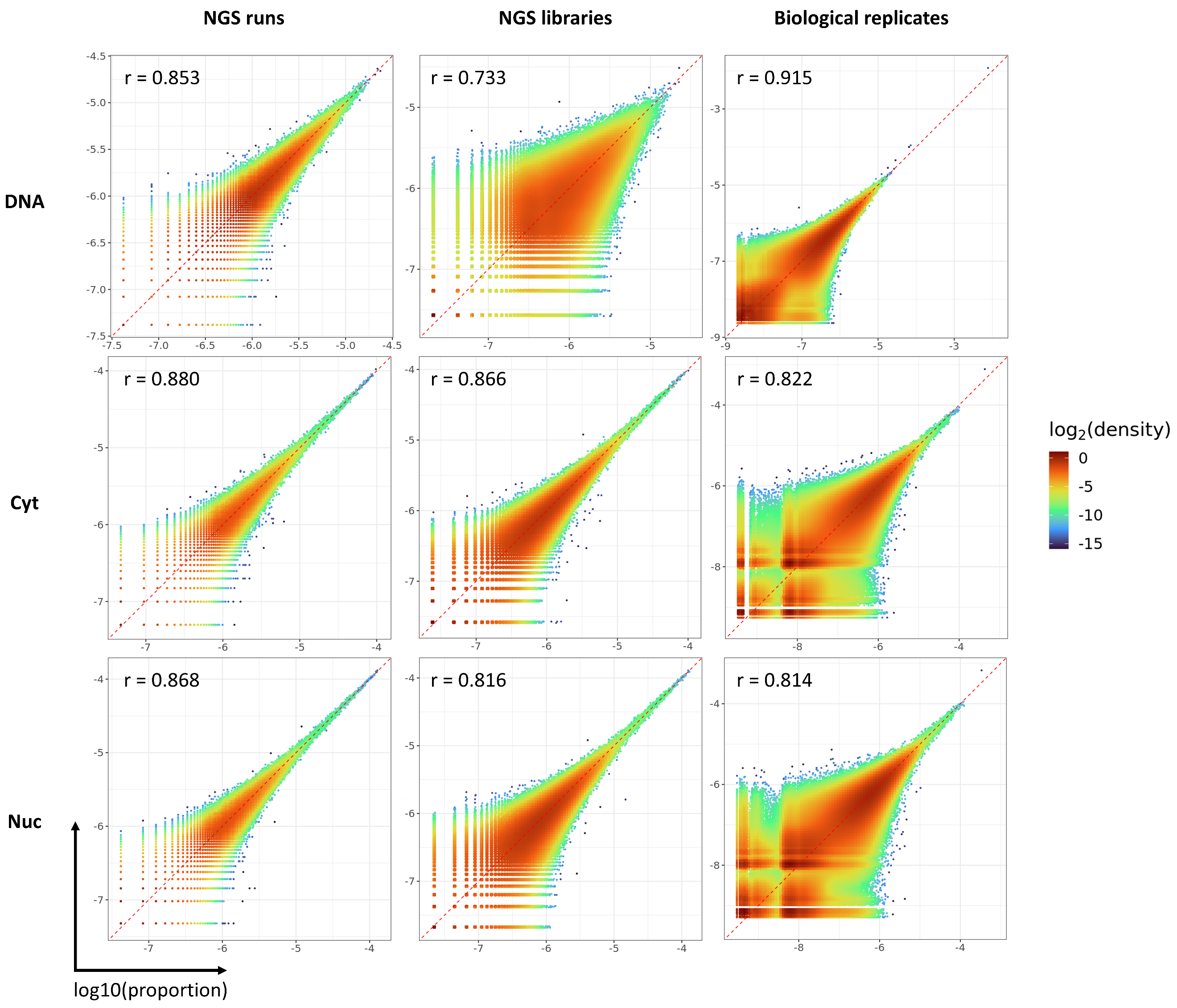

**Supplemental Fig. S3: Reproducibility of SEERS measurements in A549.** Concordance was assessed across NGS libraries generated from each fraction (DNA, cytoplasmic RNA, and nuclear RNA), including (i) repeated sequencing runs of the same library (NGS runs), (ii) independent library preparations from the same sample (NGS libraries), and (iii) independent biological replicates (biological replicates). The panels show representative examples rather than the full set of comparisons. Pearson correlation coefficients (r) are reported. Axes indicate the read proportion of each N45 measured in the corresponding library, plotted as log10(proportion).

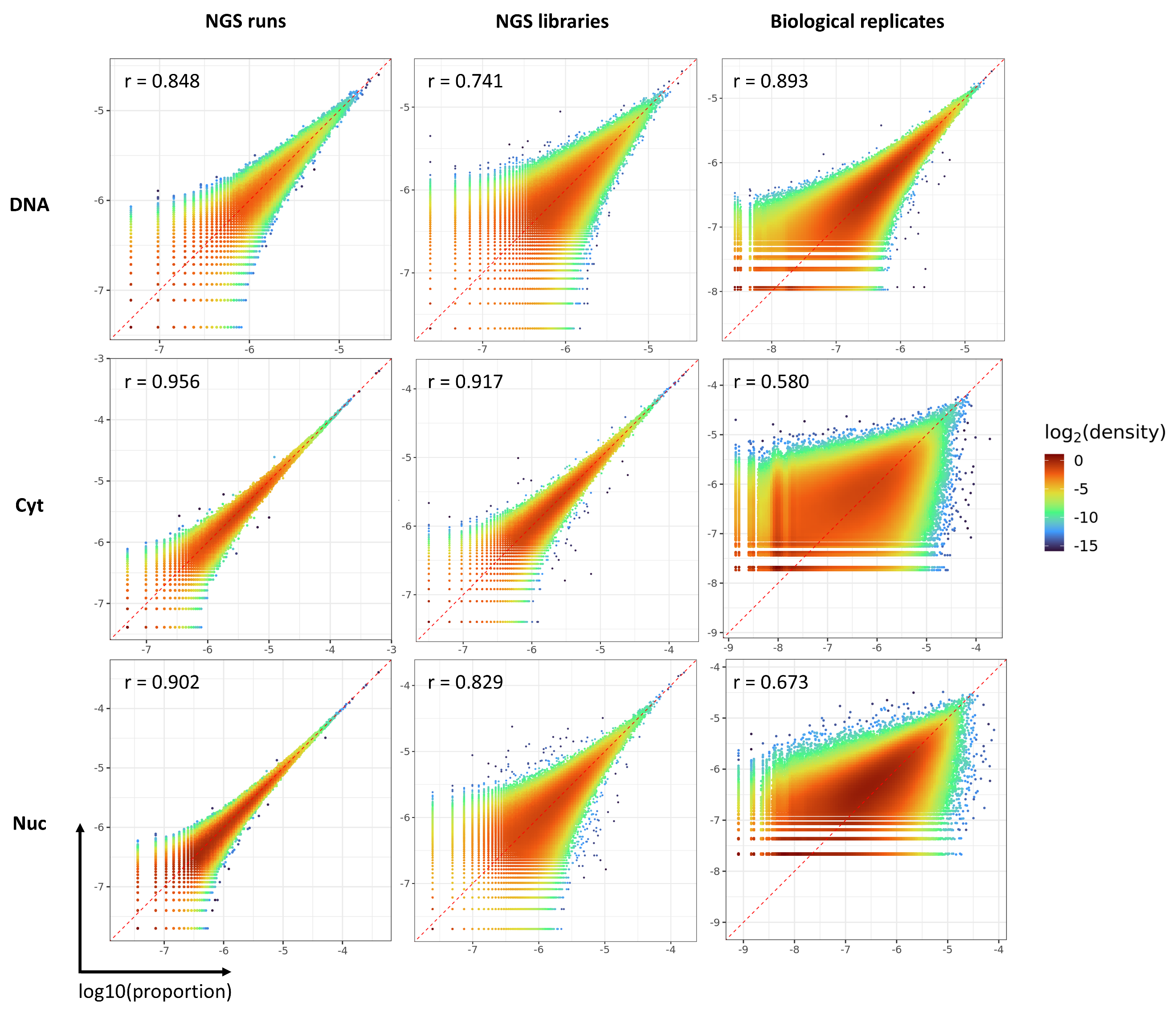

**Supplemental Fig. S4: Reproducibility of SEERS measurements in HCT116.** Concordance was assessed across NGS libraries generated from each fraction (DNA, cytoplasmic RNA, and nuclear RNA), including (i) repeated sequencing runs of the same library (NGS runs), (ii) independent library preparations from the same sample (NGS libraries), and (iii) independent biological replicates (biological replicates). The panels show representative examples rather than the full set of comparisons. Pearson correlation coefficients (r) are reported. Axes indicate the read proportion of each N45 measured in the corresponding library, plotted as log10(proportion).

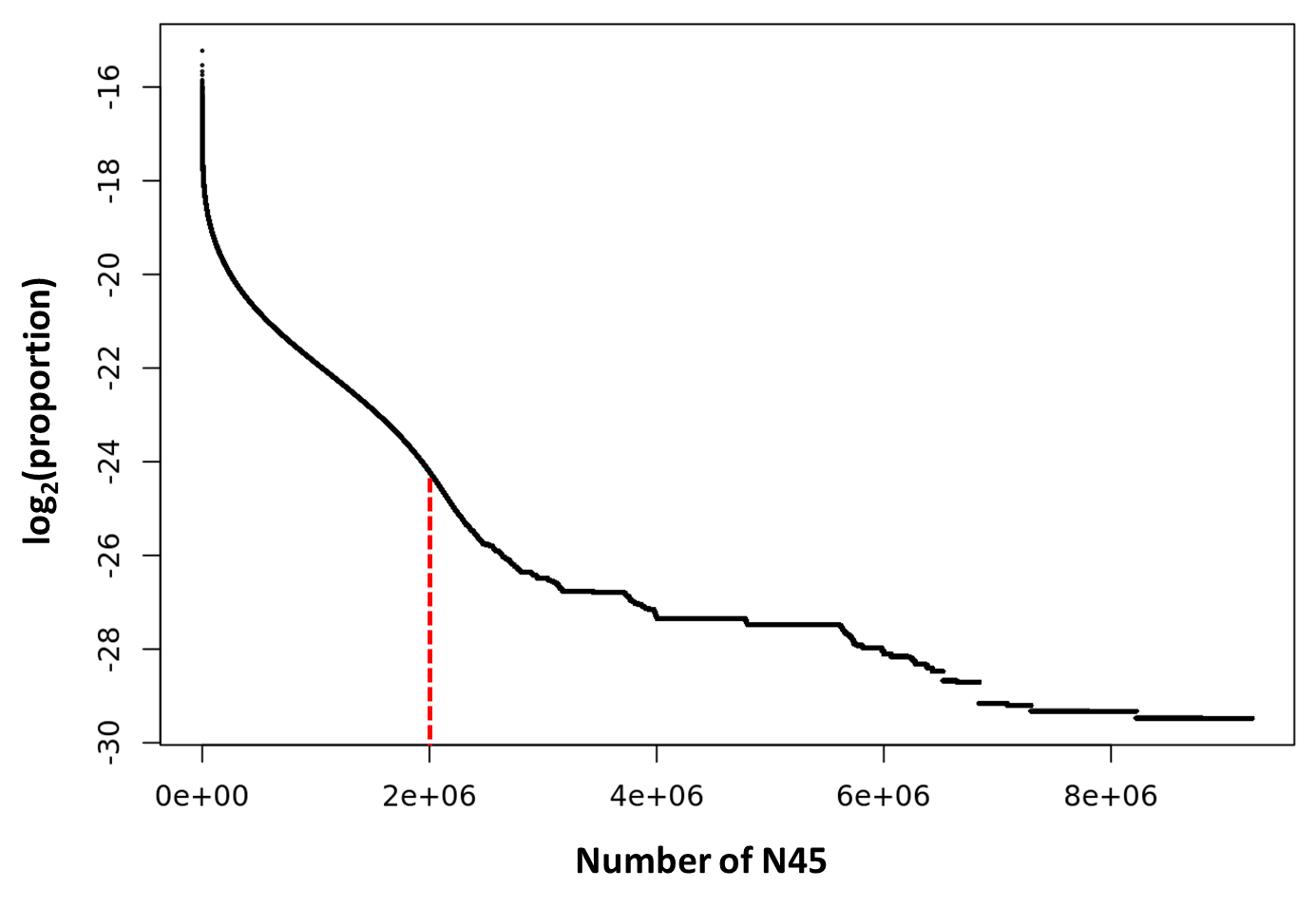

**Supplemental Fig. S5: Low-DNA-abundance N45s were removed to reduce technical noise.** Shown are the number of distinct N45 inserts detected in the DNA fraction and their relative abundances, ranked from highest to lowest. Because N45s with extremely low DNA representation are more likely to arise from sequencing errors or other stochastic technical events, they were excluded from downstream analyses. The vertical red dashed line marks the cutoff; all N45s to the right were removed, retaining ~2 million N45s for subsequent analyses.

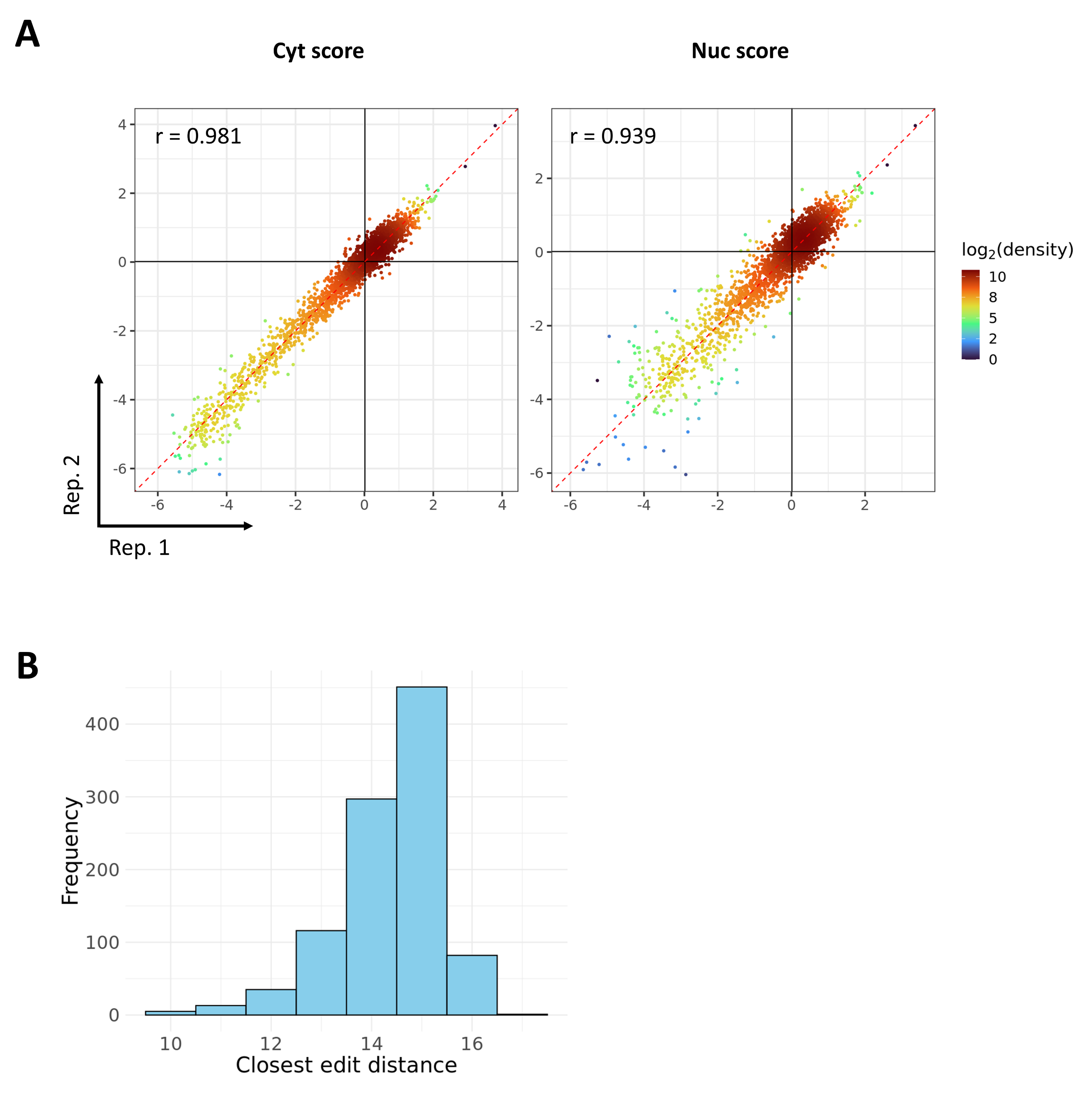

**Supplemental Fig. S6: A small-scale SEERS experiment as an independent test set.** We performed a small-scale SEERS experiment in A549 cells comprising ~3,000 N45 variants, which was used as an independent held-out dataset for benchmarking model performance. (**A**) Concordance between experimental replicates for Cyt score and Nuc score. (**B**) Distribution of the minimum edit distance between the ~3,000 N45s in this dataset and the ~2 million N45s used for model training.

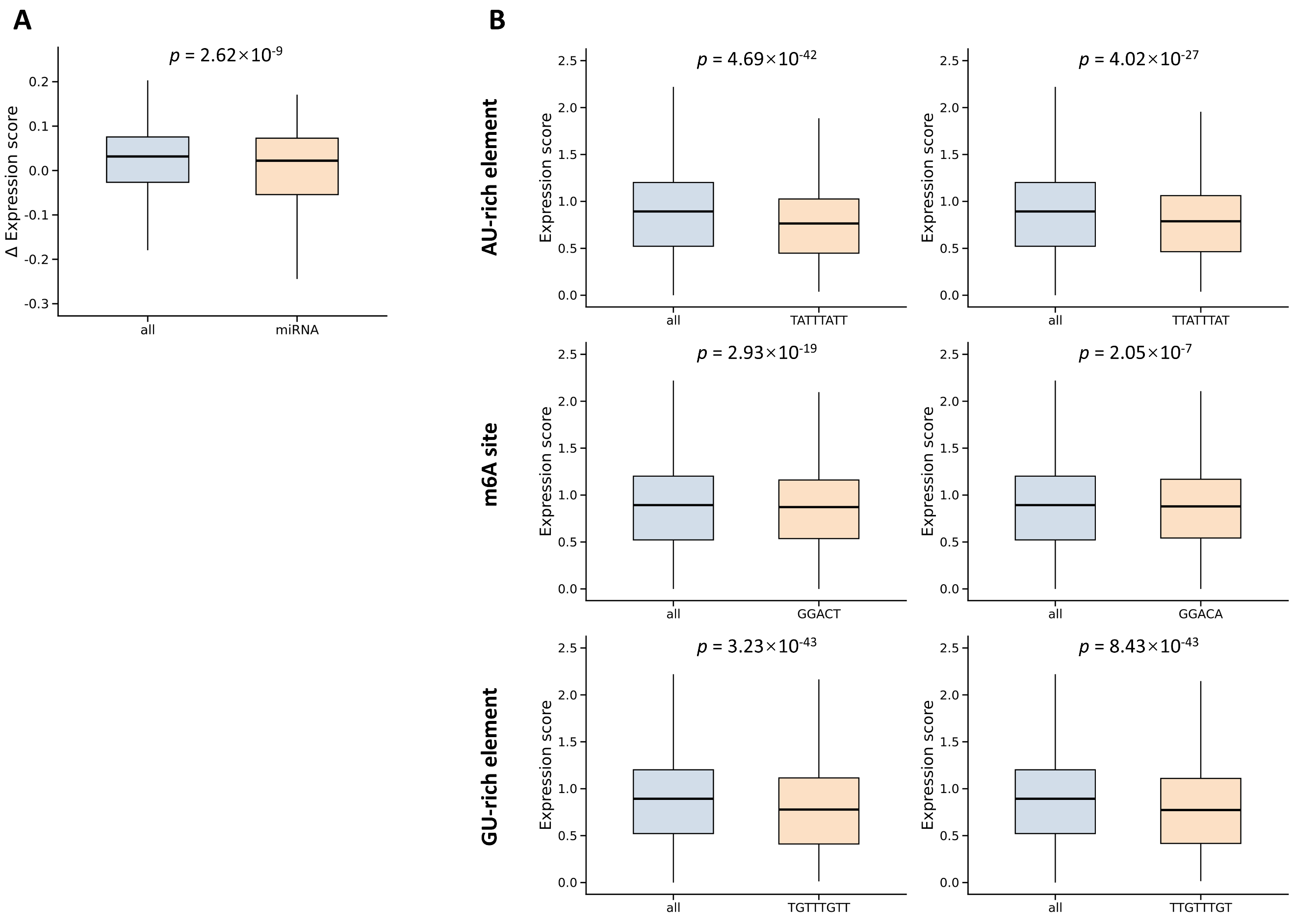

**Supplemental Fig. S7: Analysis of known 3′UTR regulatory elements in the A549 SEERS-3′UTR dataset.** (**A**) Distribution of Δ Expression scores for human miRNA binding sites (7-mers) predicted by TargetScan (n = 2,064), compared with the background set of all 7-mers (n = 16,384). p-values were calculated using Mann–Whitney U tests. (**B**) Expression score distributions for N45 inserts that contain selected regulatory-element k-mers, compared with the background distribution across all N45 inserts (n = 2,000,000). p-values were calculated using Mann–Whitney U tests.

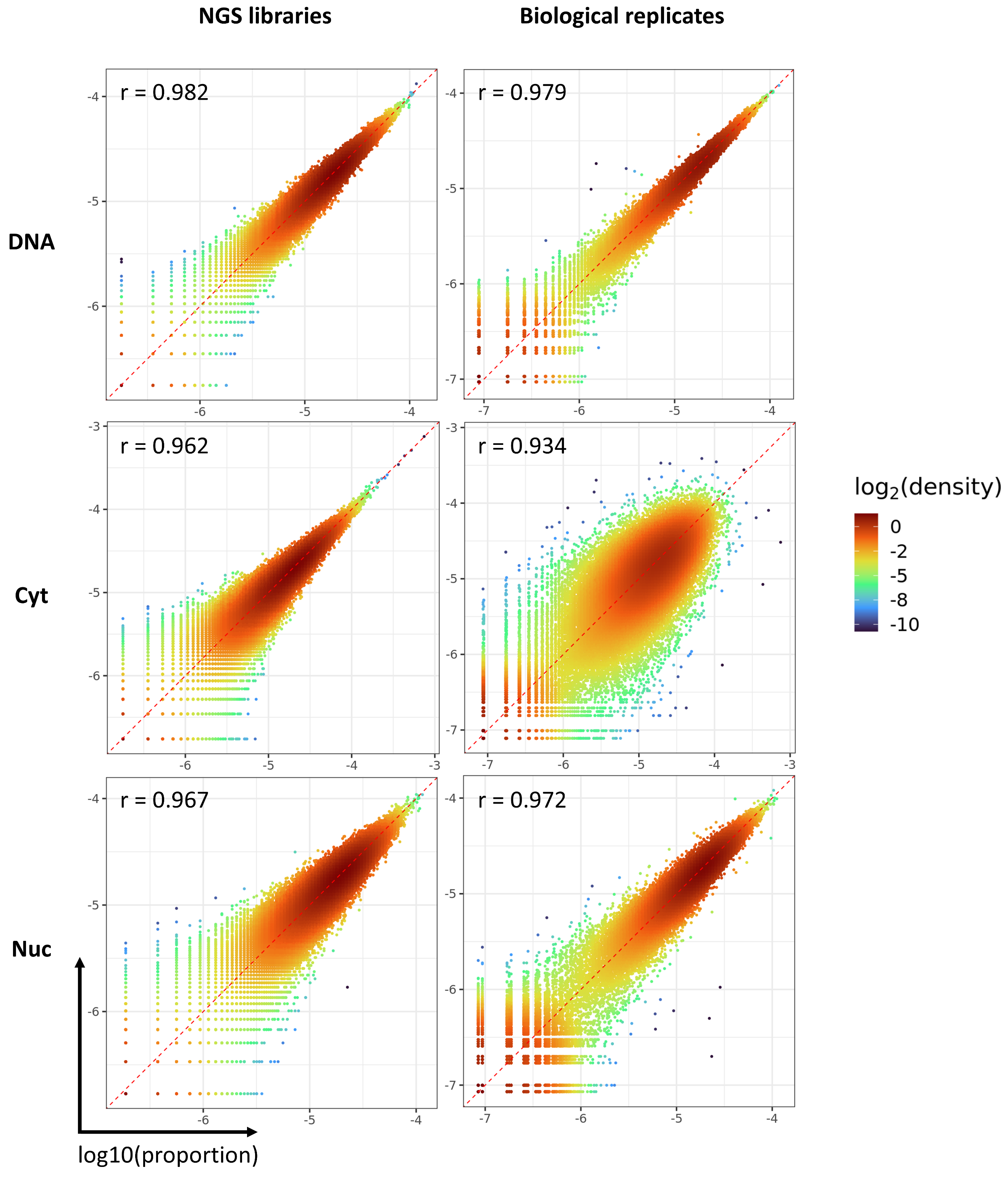

**Supplemental Fig. S8: Reproducibility of SEERS-5′UTR measurements.** Measurement concordance in the SEERS-5′UTR assay in A549 cells was evaluated across NGS libraries from three fractions (DNA, cytoplasmic RNA, and nuclear RNA). Reproducibility was assessed at two levels: (i) independent library preparations from the same sample (NGS libraries) and (ii) independent biological replicates. Panels show representative comparisons rather than the full set. Pearson correlation coefficients (r) are indicated. Axes denote the read proportion of each N45 in the corresponding library, plotted as log10(proportion).

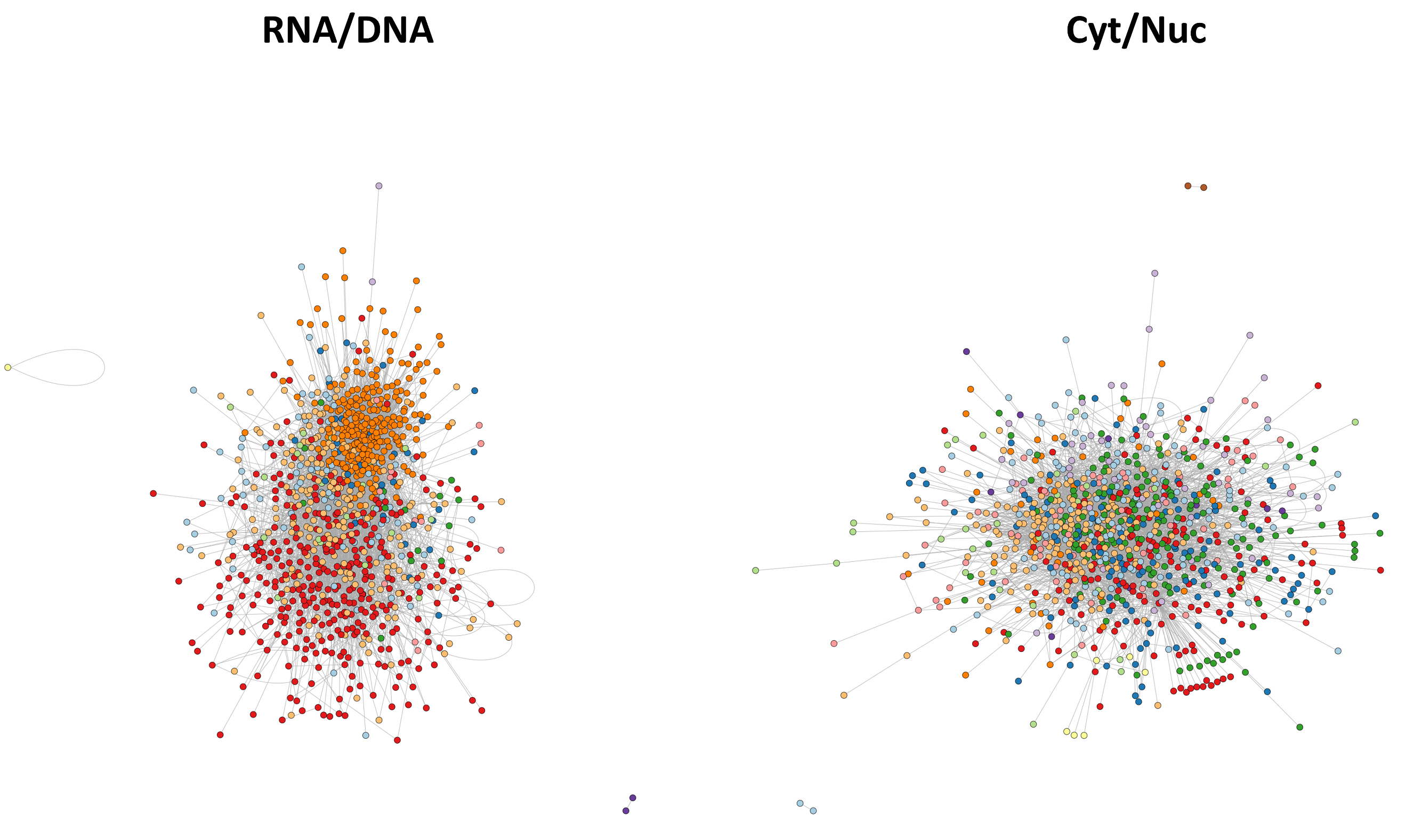

**Supplemental Fig. S9: Network of 5-mer pairs exhibiting supra-additive (synergy-like) joint effects in SEERS.** Shown are 5-mer pairs whose combined presence within the same N45 produces an effect significantly stronger than the additive expectation derived from their individual effects. Pairs are highlighted if they either increase the Expression score (enhanced transcript output) or decrease the Export score (reduced cytoplasmic enrichment).

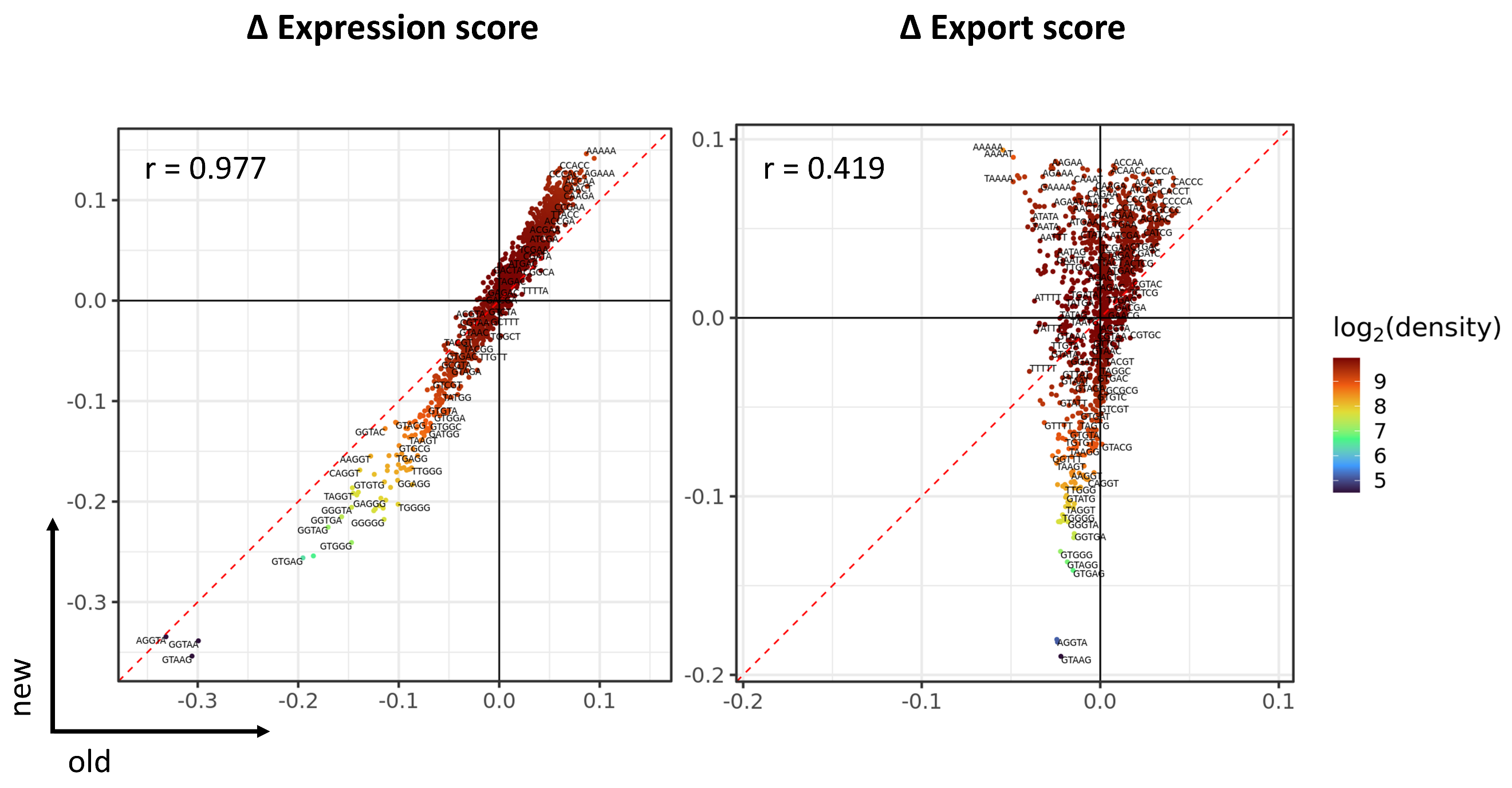

**Supplemental Fig. S10: Gene-specific priming eliminates artifactual nuclear enrichment of A-rich N45 sequences.** Comparison of 6-mer regulatory profiles in A549 derived from the original oligo(dT)-primed library preparation (old) and from the updated protocol using a gene-specific reverse-transcription primer (new). For each 6-mer under each protocol, Δ Expression score and Δ Export score were computed relative to all N45 inserts. With oligo(dT) priming, A-rich 6-mers appeared spuriously associated with reduced Export scores (apparent nuclear enrichment), whereas this pattern is abolished with gene-specific priming, indicating that the previous enrichment largely reflected library-preparation artifacts rather than genuine 3′UTR regulatory signals.
